# Enhanced muscle uptake of chemically optimized miR-23b antisense oligonucleotides as lead compounds for Myotonic Dystrophy type 1

**DOI:** 10.1101/2025.07.30.667638

**Authors:** Irene González-Martínez, Estefanía Cerro-Herreros, Marc Carrascosa-Sàez, Andrea García-Rey, Diego Piqueras Losilla, Anna Colom-Rodrigo, Nerea Moreno, Mouli Chakraborty, Aline Huguet-Lachon, Anchel González-Barriga, Neia Naldaiz-Gastesi, Martxel Dehesa, Ana Díaz-Maqueda, Nuria Barquero, Miguel A Varela, Adolfo López de Munain, Ramon Eritja, Geneviève Gourdon, Arturo López-Castel, Manuel Pérez-Alonso, Beatriz Llamusi, Rubén Artero

## Abstract

Myotonic dystrophy type 1 (DM1) is a multisystemic disorder caused by CTG repeat expansions in *DMPK*. Mutant transcripts containing expanded CUG repeats form ribonuclear foci that sequester muscleblind-like splicing regulator (MBNL) proteins, key regulators of RNA splicing and metabolism. This functional depletion leads to widespread mis-splicing and persistence of fetal transcript profiles, which underlie muscle weakness, myotonia, and muscle atrophy. In addition, miR-23b is upregulated in DM1 muscle and further represses MBNL1 translation, amplifying molecular defects. We developed chemically optimized miRNA-targeting antisense oligonucleotides (antimiRs) to inhibit miR-23b and restore functional MBNL1 levels. Using a multi-step screening process, we evaluated antimiRs with varying sequences, lengths, chemical modifications, and lipid conjugations. A key optimization was a 3’-oleic acid conjugation combined with specific chemical modifications, which enhanced muscle uptake and efficacy. Lead candidates showed strong activity in preclinical models (*HSA^LR^* and DMSXL mice, and human myoblasts), increasing MBNL1 levels, correcting mis-splicing, improving muscle strength, and reducing myotonia. They also exhibited efficient biodistribution to skeletal muscle, a critical DM1-affected tissue. *In vitro* toxicology indicated a favorable safety profile with minimal immune or renal toxicity. The antimiR mechanism was conserved in rat and pig fibroblasts. Overall, two lead antimiRs emerged as promising therapeutic candidates for DM1, with improved pharmacokinetics, tissue targeting, and safety, supporting the potential of microRNA-based approaches to correct key molecular defects in this disorder.

## INTRODUCTION

Myotonic dystrophy type 1 (DM1, OMIM#160900) is a rare genetic disorder with an overall prevalence of one in 3,000–8,000 individuals worldwide. ^1^ DM1 is characterized by expansion of an unstable CTG microsatellite repeat in the 3’-untranslated region (UTR) of *DM1 protein kinase* (*DMPK,* OMIM*605377). Mutant *DMPK* transcripts accumulate in the nucleus and induce toxicity by affecting RNA processing mechanisms, disturbing downstream regulation of gene expression.^2^ One key molecular event contributing to DM1 pathogenesis involves sequestration of the muscleblind-like (MBNL1 and 2) splicing regulator family of proteins by CUG expansions, particularly MBNL1 in skeletal muscles, which forms ribonuclear aggregates (foci).^3^ This anomalous functional depletion of MBNLs severely affects the available levels of MBNL factors, impairing their critical regulatory role in alternative pre-mRNA splicing, microRNA biogenesis, and polyadenylation.^4–6^ Additional stress responses caused by toxic *DMPK* mRNA accumulation leads to the activation of MBNL1 antagonists such as CELF1.^7^ Together, MBNL1 and CELF1 proteins function as developmental switches, and their imbalance in DM1 causes abnormal persistence of fetal patterns in processed mRNAs, resulting in inadequate protein isoforms or altered gene expression levels.^8^ This pathological mechanism affects hundreds of genes in various tissues and organs, many of which have been directly linked to the specific symptoms of DM1. ^9–12^ However, several reports support that lack of MBNL1 function contributes the most to muscular and cognitive DM1 phenotypes. ^13–19^ This is particularly favorable from a therapeutic perspective because the limited availability of MBNL1 and MBNL2 proteins, which are partially functionally redundant in the muscles,^20^ can be compensated by upregulating *MBNL1* (OMIM*606516) and/or *MBNL2* (OMIM*607327), which remain normal at the genomic level in patients. Indeed, MBNL1 overexpression effectively rescues DM1 phenotypes in preclinical models^21–23^ and is well tolerated in transgenic mice.^17^

Recent findings indicate that miR-218 and miR-23b (translational repressors of MBNL1 and 2) ^24,25^ are significantly overexpressed in diseased muscle,^25,26^ exacerbating functional protein depletion by sequestration. The observed downregulation of MBNL1 in DM1 cell models, including primary myoblasts differentiated in vitro, can be explained by elevated miRNA levels ^27^ and antagonizing these miRNAs with antimiRs increased MBNL1 protein expression and reduced *DMPK* in human DM1 cells.^27^ These observations further support miRNA inhibition as a strategy to restore MBNL1 function.

Beyond these molecular mechanisms, DM1 exhibits a highly variable clinical presentation influenced by repeat instability and multisystem involvement. ^28^ Due to errors in DNA replication and repair mechanisms, these repeats can expand during genetic transmissions, a phenomenon known as genetic anticipation, leading to earlier onset and more severe symptoms in successive generations. The length of the CTG repeat broadly correlates with disease severity, but somatic and germline instability, as well as non-CTG interruptions in expanded alleles, contribute to the variable clinical presentation of DM1.^29^ Skeletal muscle alterations, including muscle atrophy and myotonia, cause disabling symptoms. Muscle atrophy leads to progressive weakness, especially in distal muscles, impairing mobility and dexterity. Myotonia, characterized by delayed muscle relaxation, causes stiffness and difficulty in gripping or walking, further limiting physical function and independence.^1^ DM1 patients suffer from clinically relevant multisystem manifestations. In the cardiac muscles, arrhythmias and conduction defects may lead to sudden cardiac death. Progressive heart muscle degeneration, marked by fibrosis and fatty infiltration, impairs systolic and diastolic function.^30^ The smooth muscles in the gastrointestinal tract are also impacted, leading to symptoms such as dysphagia, constipation, and other digestive issues.^31^ Additionally, respiratory muscles are weakened, which contributes to breathing difficulties and increases the risk of respiratory failure.^32^ Central nervous system effects cause features such as hypersomnia, behavioral autism spectrum disorder, and cognitive deficits.^33,34^ Despite several ongoing drug development programs, ^35^ as yet no specific drug has obtained regulatory approval for DM1 patients.

AntimiRs are synthetic nucleic acid strands designed to bind to specific miRNA molecules, thereby modulating downstream gene expression with precision. Although many closely related splice-switching oligonucleotides and gapmers have reached clinical trials,^36^ no antimiRs are currently used in medical practice. Research in the medicinal chemistry of nucleic acids has experienced substantial and rapid advances aimed at improving their pharmacokinetic (PK) and pharmacodynamic (PD) properties. ^37^ For instance, nuclease resistance and enhanced plasma protein binding (leading to reduced clearance) are markedly improved by substituting the natural phosphodiester inter-nucleotide bonds with phosphorothioate (PS) linkages or incorporating locked nucleic acids (LNAs).^38,39^ Some of these features can also be achieved using nucleotides with pentose modifications, including various 2’-modified sugars such as 2’-O-methyl (2’-O-Me), 2’-O-methoxyethyl (2’-MOE), and 2ʹ-fluoro (2’-F) modifications.^37,40^ However, incorporating some of these modifications into candidate antisense oligonucleotide (ON) drugs can result in toxic accumulation in the liver and kidneys due to nonspecific interactions of the ON backbone with proteins, contributing to some degree of hepatic and renal damage.^41–43^ Moreover, antisense ON uptake by several tissues and cell types is poor, one of the most challenging being skeletal muscles, as shown in patients with the first ON that reached clinical trials in DM1.^44,45^ One active research approach to solve these issues is to conjugate the ON to an antibody^46^ or fragment of antibody for cell receptor recognition or to conjugate to small biomolecules that can influence the biodistribution of the molecule or its half-life *in vivo*.^47–49^ Therapeutic ONs represent a rapidly developing and expanding therapeutic platform for treating neuromuscular and other disorders, resulting in nearly twenty approved ON-based therapies^37,50,51^ for a variety of conditions, though none yet for DM1.

We report a streamlined approach to developing antimiR-23b hits^24,52,53^ into lead drugs for human use, by addressing chemical optimization and ON conjugation in parallel and thus significantly saving time and resources. Using the therapeutic index (T_index_) to prioritize candidates with the greatest therapeutic window, alongside established knowledge on nucleic acid medicine,^54^ top *in vitro* candidates underwent *in vivo* evaluation for dose-responsiveness, duration of activity, toxicity, immune activation, and organ biodistribution, and the mechanism of action was confirmed in rat and pig fibroblasts.

## MATERIAL AND METHODS

### Study design

The design encompasses antimiR-23b molecule optimization and testing both *in vitro* and *in vivo* to assess efficacy, toxicity, and immune activation potential, to select a lead compound with the highest potential for subsequent promotion to human use. For *in vitro* studies, antimiR candidates were screened using various chemical modifications and conjugates. The screening evaluated their EC_50_, TC_50_, and T_index_ in TDMs. The *in vivo* experiments used DMSXL and *HSA^LR^* mouse models. AntimiR candidates were administered to *HSA^LR^* mice in different doses (3, 6, 100 mg/kg), and at 25 mg/kg to DMSXL mice, after which relative muscle strength and myotonia were evaluated. Molecularly, key readouts included quantification of miRNA and MBNL1 protein and transcript levels, and alternative exon inclusion levels in skeletal muscles. Toxicity and immune activation assays were also conducted using multiple antimiR doses. RPTEC-TERT1 cells were used to assess potential toxicities and cell proliferation, while human peripheral blood mononuclear cells (PBMCs) were utilized to measure cytokine release and complement activation. Additionally, transcriptome-wide RNA-seq and Sylamer analyses were performed to assess the global impact of miR-23b inhibition on gene expression and splicing biomarkers in patient-derived muscle cells.

### Oligonucleotide synthesis

ON were provided by AXOLABS and resuspended in 1x PBS to prepare 5 mM stock solutions, stored at −20°C. 3’-oleyl-antimiRs were synthesized according to ^55^. Tables S1–S3 include the sequence, modifications, length, concentrations tested, and TC_50_, EC_50_, E_max_ and T_index_ parameters of each tested ON.

### Cell culture experimentation

Immortalized MyoD-inducible (doxycycline) DM1 and control fibroblasts were kindly provided by Dr. Furling (Institute of Myology, Paris) and were grown in Dulbecco’s Modified Eagle Medium (DMEM) with 4.5 g/l glucose, 1% penicillin/streptomycin, and 10% FBS (Sigma, Saint Louis, Missouri).^56^ Fibroblast trans-differentiation into myotubes was performed according to ^24^. Trans-differentiation was induced at day 0, and test compounds were added to the cell medium at the different concentrations indicated 24 h later by lipofection with X-tremeGENE™ HP (Roche, Basel, Switzerland) and were replaced with fresh differentiation medium 4 h afterward. Cells were collected on day 4 in the differentiation medium and processed for protein extraction.

RPTEC/TERT1 cells (CHT-003-0002, EVERCYTE GmbH) were cultured in phenol red-free DMEM/F12 medium (Gibco™ 11039021) supplemented with 1% penicillin/streptomycin, 3.5 µg/ml L-ascorbic acid (A4544), 25 ng/ml prostaglandin E1 (P5515), 8.65 ng/ml sodium selenite (S5261), 25 ng/ml hydrocortisone (H0396), 10 ng/ml human EGF (E9644), 5 µg/ml human insulin (I9278), 5 pM 3,3’,5-triiodo-L-thyronine (T6397) [all from Sigma-Aldrich], 100 µg/ml geneticin (InvivoGen, ant-gn-5), 5 µg/ml human transferrin (Merck-Millipore, 616424), and 1% fetal bovine serum (Gibco™ 10270106). Cells were maintained in 25 cm² flasks and passaged 1–2 times per week using a 1:2 to 1:3 subculturing ratio, following the manufacturer’s guidelines.

For ON toxicity assessment, RPTEC/TERT1 cells were seeded into 96-well plates (Greiner Bio-One™, 655180) at 20,000 cells/well in the specified medium and grown to confluence. On day 0, cells were treated with antimiRs prepared in PBS at final concentrations of 1, 10, 30, and 100 µM in 50 µl volumes. A toxic ON, SGLT2-MOE-ON (66), served as a positive control at 20, 300, 400, and 500 µM in 50 µl volumes. Final ON solutions were added to 150 µl of complete medium, with PBS as a vehicle control. On day 6, the medium was collected, centrifuged at 10,000 rpm to remove sediment, and the supernatants were stored at −80°C for EGF evaluation.

Primary myoblasts were grown on 0.1% gelatin-coated flasks in a proliferation medium containing DMEM supplemented with 10% FBS, 22% M-199, PSF 1x, insulin 1.74 μM, L-glutamine 2 mM, FGF 1.39 nM and EGF 0.135 mM. When cells reached 80–90% confluence, the medium was substituted by differentiation media containing DMEM supplemented with 2% FBS, 22% M-199, PSF 1x, insulin 1.74 μM and L-glutamine 2 mM. At this point, cells were transfected with X82107 at 100 nM using Ribocellin (BioCellChallenge, #RC1000) as a transfection reagent following the manufacturer’s instructions. Cell pellets were collected for DNA, RNA, and protein analyses 5 days after differentiation started when treatments were also transfected.

Rat embryo (*Rat2*, ATCC CRL-1764) and pig testis (ST, ATCC CRL-1746) fibroblasts were grown in a complete medium (the same as for human fibroblasts) and seeded at 10^5^ cells/ml in petri plates. These cells were treated with X82108 at 50 nM and transfected with the same procedure as DM1 and control fibroblasts.

### RNA extraction, semi-quantitative RT-PCR and RT-qPCR

Total RNA from murine gt and qd muscle was isolated using the miRNeasy Mini Kit (Qiagen, Hilden, Germany) according to the manufacturer’s instructions. One microgram of RNA was digested with DNase I and reverse-transcribed with SuperScript II (Invitrogen, Carlsbad, California) using random hexanucleotides. For subsequent PCR reactions, 20 ng of cDNA was used with GoTaq polymerase (Promega, Madison, Wisconsin). Specific primers were used to analyze the alternative splicing of *Atp2a1* (OMIM*108730), *Nfix* (OMIM*164005), *Mbnl1*, and *Clcn1* (OMIM*118425) in mouse samples (both muscles). *Gapdh* (OMIM*138400) levels established the endogenous reference using 0.2 ng of cDNA. PCR products were separated on a 2% agarose gel and quantified using ImageJ software (NIH, Bethesda, Maryland). The PSR index per alternative exon was defined as the value: %EI minus X̅%DEI, divided by X̅%DEI minus X̅%HEI (EI: exon inclusion of each sample; DEI: disease exon inclusion; HEI: healthy exon inclusion). Overall PSR was the average of the above PSR for the four quantified alternative splicing events and the two muscles analyzed. The primer sequences and exons analyzed are available in ^24^.

We used 1 ng of mouse tissue cDNA as a template for multiplex RT-qPCR using the QuantiFast Probe PCR Kit reagent. Commercial TaqMan probes (Qiagen, Hilden, Germany) were used for mouse (*Mbnl1* and *Mbnl2*; FAM-labeled probes) and reference (*Gapdh* MAX-labeled probe) genes. Results were normalized to *Gapdh* endogenous expression. The sequences of the probes and primers are included in Table S4.

MiRNA expression in muscle tissues was quantified using specific miRCURY™-locked nucleic acid microRNA PCR primers (Qiagen, Hilden, Germany) according to the manufacturer’s instructions. Relative gene expression was normalized to U1 (YP00203909) and U6 (YP00203907) snRNAs. Expression levels were measured using a QuantStudio 5 Real-Time PCR System (Applied Biosystems, Foster City, California). Expression relative to the endogenous gene and control group was calculated using the 2^−ΔΔCt^ method.

### Animal experimentation and antimiR administration

Mouse handling and experimental procedures for *HSA^LR^* and FVB animals followed the European law regarding laboratory animal care and experimentation (2003/65/C.E.) and were approved by the regional Ministry for Agriculture (Conselleria de Agricultura, Generalitat Valenciana). Homozygous transgenic *HSA^LR^* (line 20 b) mice^57^ were provided by Prof. C. Thornton (University of Rochester Medical Center, Rochester, NY, USA). Experimental groups were FVB as wild-type control, *HSA^LR^* treated with PBS as negative control, and *HSA^LR^* mice treated with each experimental antimiR. The sample size was four mice per treatment group, twelve mice for PBS, and eighteen mice for the FVB group. All groups were injected intravenously (tail vein) with 1×PBS (vehicle) or the specific antimiR with a single dose of 3 or 6 mg/kg, except for the 100 mg/kg group, which received a single subcutaneous injection. At 5, 7 or 15 days after injection, the mice were sacrificed, and the tissues of interest were frozen in liquid nitrogen for molecular assays. Additionally, DMSXL mice (>99% C57BL/6 background) provided by Prof. G Gourdon (Sorbonne University, Paris, France), carrying over 1500 CTG repeats within the *DMPK* transgene,^58,59^ were used to study the effects of antimiR treatment (authorization APAFIS#30275-2021030414389550). These mice were 3 weeks old at the start of the experiment and received subcutaneous injections of antimiR X821 at a dose of 25 mg/kg, administered twice per week for 4 weeks. Age-matched wild-type littermate mice served as controls. Following the treatment regimen, the gastrocnemius and tibialis muscles were collected, rapidly frozen in liquid nitrogen, and stored at −80°C for subsequent molecular analyses.

### Electromyography studies

Electromyography was performed before treatment and at the time of sacrifice under general anesthesia, as previously described.^22^ Myotonia was assessed from a total of 10 needle insertions (5 per quadriceps). The overall myotonia score obtained from these insertions was then normalized to a 5-point scale, defined as follows: 0, no myotonia; 1, occasional myotonic discharges present in ≤50% of the insertions; 2, myotonic discharges present in >50% of the insertions; 3, myotonic discharges in nearly all insertions; and 4, myotonic discharges in all insertions.

### Forelimb grip strength test

Forelimb grip strength was measured using a grip strength meter (BIO-GS3; Bioseb, Pinellas Park, Florida). Peak pull strength (g) was recorded on a digital strength transducer as the mouse released its grip. The transducer was reset to 0 g after each measurement, and three measurements were taken at 30 s intervals. Body weight was recorded simultaneously, and normalized grip strength was calculated by dividing average strength by body weight. Experiments were conducted with coded animals to prevent bias.

### ELISA determinations

Custom hybridization-based ELISA determinations of antimiR conjugate concentrations in gt and qd from treated *HSA^LR^* mice were performed as described in ^60^ using the following PS probe: miR-23b (5’->3’) [BIO] ATCACATT_o_G_o_C_o_C_o_A_o_G_o_GGATTACC [DIG] double-labeled with digoxigenin (DIG) and biotin (BIO). A control sample (*HSA^LR^* treated with PBS) was used to subtract the background.

### Extracellular EGF assay

To determine extracellular EGF, Human EGF ELISA Kits (Invitrogen™ KHG0062) were used, and EGF concentration was determined according to manufacturer’s instructions. The experimental protocol was adapted from ^61^.

### *In vitro* toxicity assay

The French Blood Donors Organization (EFS, Etablissement Français du Sang) provided human serum from healthy volunteers. Complement activation, coagulation, and platelet activation were performed by SQY Therapeutics (France) in human blood samples. Test compounds were incubated with human serum, and complement activation was evaluated using commercial ELISA measurement of human C3a. The extrinsic and intrinsic coagulation pathways were evaluated in human plasma incubated with test compounds. PT and aPTT were measured on a semi-automated START max coagulometer (Stago). Flow cytometry assays evaluated human platelet activation by test compounds by detecting surface antigens (usually glycoproteins) expressed during platelet activation. Assays with PBMCs were performed by Axolabs GmbH (Germany).

### RNAseq assay

Total RNA was extracted from DM1 and control immortalized MyoD-converted fibroblasts ^56^ 72 h after transfection with 200 nM X82108 using X-tremeGENE HP (Roche) following culture in DMEM GlutaMAX with 10% FBS. RNA was purified with the RNeasy Mini Kit (QIAGEN). RNA sequencing was performed by Novogene using Illumina platforms. Poly(A)+ RNA was isolated using oligo-dT magnetic beads, fragmented, and converted to first- and second-strand cDNA. Libraries were prepared following standard end repair, A-tailing, adaptor ligation, size selection, and PCR amplification, and were assessed by Qubit, qPCR, and Bioanalyzer before pooling and sequencing. Paired-end reads were aligned to the human reference genome (GRCh38.p14) using STAR (v2.7.11b) with splicing-aware parameters and GeneCounts enabled. BAM files were processed with SAMtools, and gene-level quantification was performed with featureCounts.

Sylamer (v18-131) was used to assess enrichment of the miR-23b seed motif (AATGTGA) across differentially expressed genes (adjusted p < 0.05), ranked by log₂ fold change. Corresponding 3′UTR sequences were retrieved from hg38 using TxDb.Hsapiens.UCSC.hg38.knownGene and BSgenome.Hsapiens.UCSC.hg38, ordered according to gene ranking - with positive values corresponding to upregulated genes and negative values to downregulated genes-, and analyzed using a window growth parameter of 10. Output tables were used to visualize motif representation across bins and to overlay statistical thresholds (nominal ±p = 0.05 and ±E = 0.01).

Alternative splicing was quantified using rMATS (v4.1.2) with 150-nt unstranded read settings. Analyses were restricted to a validated panel of 22 DM1-relevant splicing events described in Provenzano *et al*.^62^ Percent Spliced-In (PSI) values were calculated as the fraction of exon inclusion isoform reads relative to all exclusion and inclusion isoform counts. Percent Splicing Rescue (PSR) for each replicate x and event y was computed as: (PSIx, y – PSIMedian Control, y)/(PSIMedian DM1, y – PSIMedian Control, y).

RNA-seq data are publicly available with accession number S-BSST2306 in the BioStudies database.

### Statistical analyses

TC_50_ values were calculated using non-linear least-squares regression. For molecular and functional parameters, one-way ANOVA was performed where applicable, preceded by the Shapiro-Wilk normality test. Normal samples were Brown-Forsythe corrected if required, while non-normal samples were analyzed using Kruskal-Wallis without Dunn correction. Differences relative to control samples were assessed with two-tailed Student’s t-test (α = 0.05), applying Welch’s correction as needed. Sample sizes (n) are indicated in each figure.

The values obtained for the *HSA^LR^* mice are represented (Figure 4C) as the recovery index (RI), which measures the proximity of the different parameter values obtained with treated *HSA^LR^* mice to those of FVB controls. This RI was obtained for the different parameters (MBNL1 protein, *Mbnl1 and Mbnl2* and miR-23b or −218 expression level, PSR, *Mbnl1* ex5 inclusion recovery, and functional recovery) of each mouse after treatment according to the following formula: value % MT minus X% MNT, divided by X% MH minus X% MNT [where MT is the value of each mice treated (PBS or oligonucleotide), MNT is *HSA^LR^* mice treated with PBS, and MH is healthy mice value (FVB)]. These values range from 0 to 100, where 0 are untreated mice (H*SA^LR^*-PBS) and 100 are healthy mice (FVB).

## RESULTS

### *In vitro* screening of antimiR-23b ON

In our search for the most effective and safest antimiR-23b, we performed *in vitro* screenings in DM1 fibroblasts transdifferentiated into myotubes (TDMs). The first screening addressed specific miR-23b-targeting sequences, length, and chemical modifications. In TDMs transfected with each candidate antimiR variant, we established the T_index_ as E_max_* (TC_50_/EC_50_), which integrates the maximum activity (E_max_), median toxic concentration (TC_50_), and median effective concentration (EC_50_) into a single metric. EC_50_ and E_max_ values were obtained through a Quantitative Dot Blot assay^63^ for MBNL1 protein quantification to measure antimiR activity, and TC_50_ was determined using an MTT/MTS toxicity assay. In all cases, we included for comparison purposes the antimiR-23b molecule used in our previous proof-of-concept studies, here called X820,^26,52^ with the sequence 5’-GGUAAUCCCUGGCAAUGUGAU-3’, 2’-OMe chemistry, PS linkages at end-proximal nucleotides, and conjugated to cholesterol at the 3’ end, and its unconjugated version (X82--).

A panel of 21 unconjugated antimiR-23b molecules containing distinct sequence and chemistry variants were synthesized and tested for their T_index_ *in vitro* (Figure 1A,B and Table S1). The results identified X821 as the most promising antimiR-23b candidate, with a T_index_ of 4018.3. This ON demonstrated a 423-fold improvement compared to X82-- (T_index_: 9.5), our unconjugated benchmark, and a 5.9-fold improvement over the second-ranked X822 (T_index_: 686.7). X821 exhibited the best toxicity profile *in vitro*, requiring around 16 times higher concentration than X820 to reach TC_50_ (Figure 1B and Table S1). Comparing unconjugated versions among themselves, 18 of the 21 antimiRs evaluated had a T_index_ higher than X82--, while three worsened the parameter.

**Figure 1.**
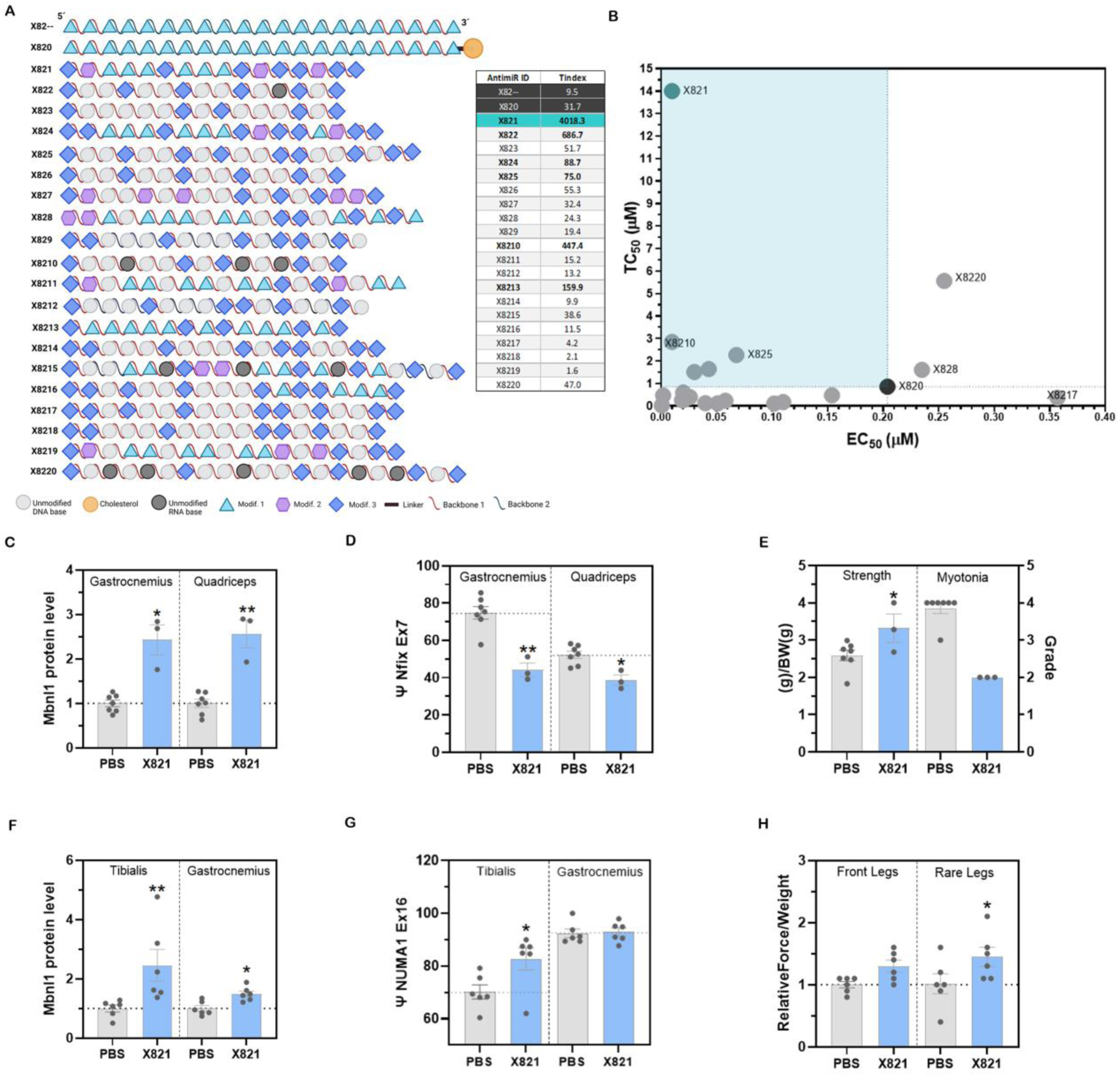
*In vitro* screening of antimiR-23b variants and assessment of safety and efficacy of a prospective lead candidate in two mouse models. **(A)** Schematic representation of the antimiRs tested in the screening. Modifications 1, 2, and 3 refer to 2’-O-Methyl RNA nucleotides, 2’-O-MOE RNA nucleotides, and LNA nucleotides. Backbones 1 and 2 represent a mix of PS (phosphorothioate) and PO (phosphodiester) linkages. **(B)** Representative graphs illustrating the ranking of the compounds tested in the screening, based on their EC_50_ (µM) and TC_50_ (µM) values. The graph also includes a table displaying the T_index_ of each antimiR. In both graph and table, the benchmark compounds are indicated in dark gray, the antimiR hit of the screening in greenish-blue, and the remaining antimiRs in gray. The parameters measured in the two mouse models (*HSA^LR^* (**C,D,E**) and DMSXL (**F,G,H**)) treated subcutaneously with X821 at SD 100 mg/kg and MD 25 mg/kg, respectively, were: **(C,F)** MBNL1 levels of treated mice relative to saline controls in skeletal muscles affected in DM1; **(D,G)** the percentage of exon inclusion (Ψ) of the indicated exons and muscles. **(E,H)** Functional rescues after treatment: **(E)** relative force and myotonia grade change compared to controls; **(H)** change in relative muscle strength of antimiR-treated mice compared to saline controls at the indicated days after treatment. Error bars indicate mean ± SEM. *: p<0.05; **: p<0.01; ***: p<0.001. The data were analyzed using unpaired Student’s t-test, compared with untreated DM1 mice (phosphate-buffered saline, PBS). AI: after injection; SD: single dose, MD: multiple dose. *HSA^LR^*: PBS n=7, X821 n=3; DMSXL: PBS n=6, X821 n=6.

### Effects of X821 on MBNL1 levels, alternative splicing, and functional muscle defects in two DM1 mouse models

Prospective lead candidate X821 was evaluated *in vivo* subcutaneously to assess its therapeutic activity in a well-established adult-onset DM1 murine model, the *HSA^LR^* mouse, which expresses expanded 250 CUG repeat tracts in human skeletal actin (HSA) transcripts. ^57^ Additionally, we examined the effectiveness of the antimiR approach in DMSXL mice, which carry a complete *DMPK* with more than 1,500 CTG repeats, providing a contrast to the muscle-specific HSA^LR^ strain and enabling evaluation of antimiR activity in the context of a full-length human transgene. ^58^ These complementary models allow for a comprehensive evaluation of the therapeutic efficacy of X821.

In the *HSA^LR^* mouse, treatment with 100 mg/kg of X821 significantly increased MBNL1 protein levels in muscle without altering blood parameters (Figure S1A,B). Regarding C.P.K. levels, although they appear higher in the treated group, the difference compared to non-treated animals is not statistically significant. It also rescued *Nfix* ex 7, *Atp2a1* ex 22, and *Clcn1* ex 7a mis-splicing in both gastrocnemius (gt) and quadriceps (qd) muscles (Figure 1C,D, S1C,D), leading ultimately to a significantly reduced myotonia grade and increased grip strength in treated mice (Figure 1E). In the DMSXL mouse, X821 also increased MBNL1 protein levels in the tibialis anterior (TA) and gt muscles and rescued *Numa1* (OMIM*164009) exon 16 inclusion^64^ in TA (Figure 1F,G). Molecular rescues were sufficient to produce a significant functional improvement in grip strength in the rear legs 2 weeks after the last injection (Figure 1H). Taken together, these findings demonstrate that X821 effectively addressed key molecular defects, such as MBNL1 protein upregulation and alternative splicing correction in both mice, while also improving functional outcomes related to myotonia and muscle strength. Even though X821 effectively targeted miR-23b, the effects observed on muscle function did not surpass those achieved with X820 at a lower concentration.^52^ This lack of correlation between *in vitro* and *in vivo* activity underscored the need for subsequent optimization of X821 to enhance its therapeutic potential.

### *In vitro* screening of lipidic conjugates to improve the therapeutic index

A second *in vitro* screening addressed conjugation to various moieties to improve potency and PK properties.^65,66^ Conjugating hydrophobic molecules to antisense ON and employing various strategies to transport them from blood to tissue might improve their transport to muscle tissues, leading to enhanced biodistribution and functional uptake in muscle cells. ^67,68^ Fatty acid conjugation, as yet not attempted with anti-miR-type molecules, was tested here to assess the potential effects of various ligands. Additional antimiR molecules were generated by synthetically conjugating different hydrophobic moieties (Figure 2A,B, and Table S2). To isolate the impact of the moiety, the nucleotide sequence was kept identical to the original X82--, and experiments included this unconjugated version. In this screening evaluating conjugates in the 5’ end of the ONs, oleic acid (oleyl moiety; Ol) conjugation ranked first (X8200; T_index_= 1101.4), followed by the linoleyl-antimiR (X8201; T_index_= 400.9). These two conjugates had the greatest ability to reduce cell toxicity (Figure 2B), respectively showing a 16.10- and 5.84-fold superiority to the original X820 hit, and 155.13- and 56.35-fold enhancement compared to the naked version of the antimiR. Other lipid conjugations also improved T_index_ compared to the unconjugated benchmark X82-- (tocopherol, X8202; cholesterol, X8203), while conjugation to palmitic, elaidic, and stearic acids was similar or lower in terms of T_index_.

**Figure 2.**
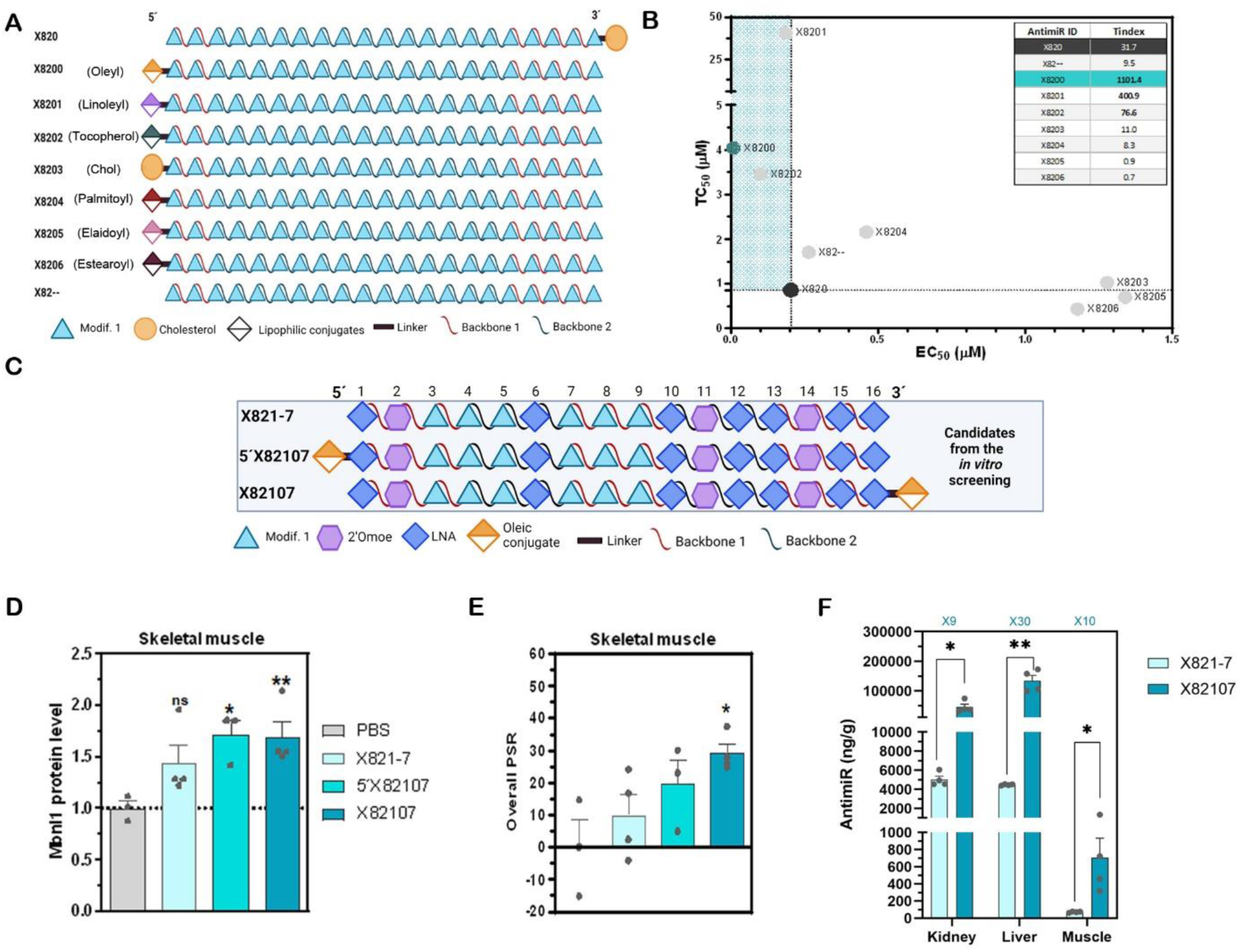
*In vitro* screening of antimiRs-23b with different conjugates and assessment of safety and efficacy of a group of lead candidates in *HSA^LR^* mice. (**A**) Schematic representation of the antimiRs tested in the screening. Modifications 1, 2, and 3 refer to 2’-O-Methyl RNA nucleotides, 2’-O-MOE RNA nucleotides, and LNA nucleotides. Backbones 1 and 2 represent a mix of PS (phosphorothioate) and PO (phosphodiester) linkages. (**B**) Representative graph illustrating the ranking of the compounds tested based on their EC_50_ (µM) and TC_50_ (µM) values. The graph also includes a table displaying the T_index_ of each antimiR. In both graph and table, the benchmark compounds are indicated in dark gray, the antimiR hit of the screening in greenish-blue, and the remaining antimiRs in gray. **(C-F)** Selected compounds were tested *in vivo* using a single 3 mg/kg IV injection in *HSA^LR^*mice. After 5 days, several parameters were measured, and skeletal muscle tissues were dissected for analysis. **(C)** Schematic representation of the antimiRs tested *in vivo*. **(D)** MBNL1 levels of treated mice relative to saline controls in skeletal muscle 5 days after injection (mean value of gt and qd). **(E)** Overall percentage splice recovery (PSR) of leading antimiR candidates in both muscles. **(F)** AntimiR X82107 vs X821-7 levels in muscle tissues, kidney and liver detected by ELISA. The multiplier value indicates the fold-change between the different versions tested. Error bars indicate mean ± SEM. *: p<0.05; ***: p<0.001. The data were analyzed by one-way ANOVA or Kruskal-Wallis test if required, compared with untreated *HSA^LR^* mice (PBS). Individual values are indicated as data points. PBS n=3, X821-7 n=4, 5’X82107 n=3, X82107 n=4.

### *In vivo* screening of antimiR lead candidates

We hypothesized that the positive impact of Ol on T_index_ could enhance the high performance of antimiR X821, aiming for synergistic or additive effects. Two X821 variants (X821-7 and X821-8) were designed, retaining the same sequence but including known chemical modifications to improve *in vivo* druggability, such as methylated cytosines designed to enhance stability (Table S3), improve RNA target interaction, and reduce immunogenicity *in vivo* ^69^. Both X821 variants differed only slightly in terms of LNA/MOE content, with X821-8 having a higher Tm than X821-7. X821-7 served as a benchmark to evaluate the effect of Ol conjugation on *in vivo* rescue performance in *HSA^LR^* mice. Therapeutic potential was assessed by comparing the rescue of key DM1 molecular and functional parameters after a single low-dose (3 mg/kg) IV injection of unconjugated X821-7 or ON with Ol at the 5’ (5’X82107) or 3’ (X82107) ends (Figure 2C).

At 5 days after injection, muscle tissue analysis revealed that both X82107 conjugate variants consistently increased MBNL1 protein levels by 1.5-fold compared to untreated animals, unlike unconjugated X821-7 (Figure 2D). For X82107, this translated into a significant rise in the overall percentage splicing recovery (PSR; Figure 2E and S2A), generated from a selected panel of four mis-spliced genes in this model (*Atp2a1, Clcn1, Nfix,* and *Mbnl1*). Increases at the *Mbnl1* and *Mbnl2* mRNA level were significant and similar for all three antimiR variants, whereas the reduction in detected miR-23b was robust for X82107 (Figure S2B-D). ELISA determinations revealed that the oleic-conjugated antimiR X82107 achieved higher concentrations in the skeletal muscles, which are tissues relatively resistant to ON uptake, compared to the unconjugated version (Figure 2F). Also, the levels detected in the kidney and liver were higher with X82107 compared to the naked version, as was expected for this type of compound, as normally they are excreted by these organs. Functionally, normalized muscle strength improved significantly and similarly for all three antimiR variants, while myotonia reduction was higher for X82107 (Figure S2E-G). Taken together, the Ol-conjugated versions at either end of the molecule were more effective than the unconjugated version.

Similar to X821-7, we generated unconjugated X821-8, featuring a different set of chemical modifications (Table S3) and contrasted the effects of Ol (X82108) and palmitoyl [Pal] (X82148)-conjugation at the 3’-end of the molecule (Figure 3A), using Pal as a benchmark conjugate for its reported action as an effective enhancer of ON effects in muscle.^67^ AntimiRs were administered as before, and in a similar way to X821-7, both X82108 and X82148 conjugates outperformed the unconjugated version, and in comparison to Pal, Ol moiety notably increased MBNL1 protein levels, significantly improving overall PSR in skeletal muscles (Figure 3B,C; S3A). The increase at the *Mbnl1* mRNA level and reduction in detected miR-23b were significant and similar for all three X82108 variants, whereas *Mbnl2* transcript levels significantly increased only after X82108 treatment (Figure S3B-D). Accordingly, ELISA determinations revealed that X82108 achieved higher concentrations in the skeletal muscle than the unconjugated version (Figure 3D). Functionally, X82108-treated mice showed a significant increase in normalized strength and a robust reduction in myotonia grade, which dropped from 3 to less than 1 (Figure S3E–G). Overall, from these *in vivo* experiments, X82108 and X82107 were selected as new prospective lead candidates for further assessment of preclinical features.

**Figure 3.**
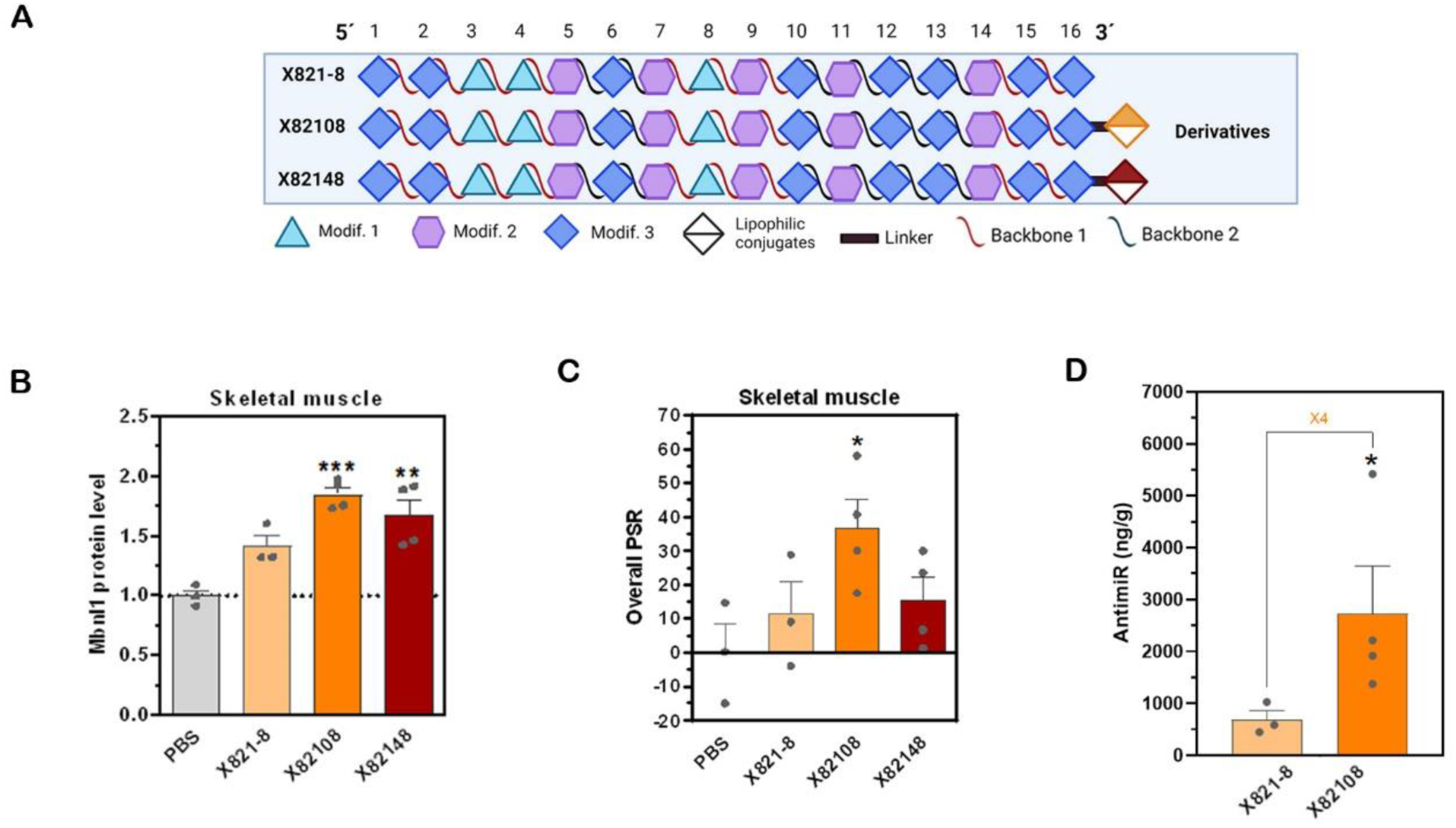
Single-dose experiment with leading antimiR derivatives from the previous experiment. **(A)** Schematic representation of the antimiRs tested. Modifications 1, 2, and 3 refer to 2’-O-Methyl RNA nucleotides, 2’-O-MOE RNA nucleotides, and LNA nucleotides. Backbones 1 and 2 represent a mix of PS (phosphorothioate) and PO (phosphodiester) linkages. Selected compounds were tested *in vivo* using a single 3 mg/kg IV injection in *HSA^LR^* mice. **(B)** MBNL1 levels of treated mice relative to saline controls in skeletal muscle (mean value of gt and qd). **(C)** Overall PSR of leading antimiR candidates in both muscles. **(D)** AntimiR X82108 and X821-8 levels in muscle tissues (mean value of qd and gt) detected by ELISA. The multiplier value indicates the fold-change between the different versions tested. Error bars indicate mean ± SEM. *: *p*<0.05; **: *p*<0.01; ****: *p*<0.0001. The data were analyzed by one-way ANOVA or Kruskal-Wallis test if required, compared with untreated *HSA^LR^* mice (PBS). Individual values are indicated as data points. PBS n=3, X821-8 n=3, X82108 n=4, X82148 n=4.

**Figure 4.**
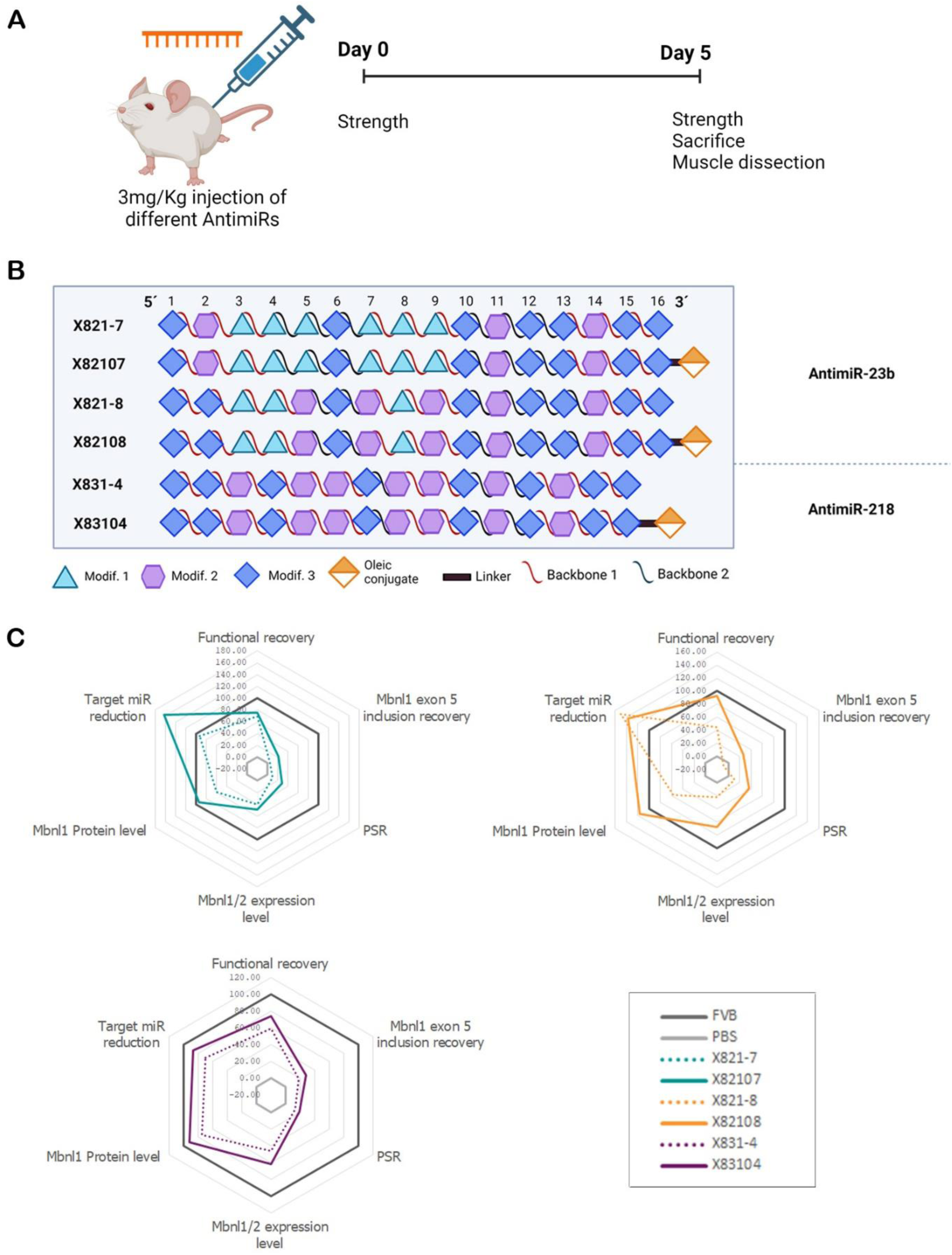
Effect of Oleyl conjugate in different antimiRs. **(A)** Schematic representation of the experimental design using a single 3 mg/kg IV injection in *HSA^LR^* mice. Modifications 1, 2, and 3 refer to 2’-O-Methyl RNA nucleotides, 2’-O-MOE RNA nucleotides, and LNA nucleotides. Backbones 1 and 2 represent a mix of PS (phosphorothioate) and PO (phosphodiester) linkages. **(B)** Schematic representation of the antimiRs tested. **(C)** Spider graphs depicting the readouts obtained in the *HSA^LR^* mouse model with leading antimiR candidates. Functional recovery includes strength and myotonia measurements. PSR includes rescues of *Atp2a1* ex22, *Clcn1* ex7a, *Nfix* ex7 and *Mbnl1* ex5 in qd and gt. FVB n=5, PBS n=3, X821-7 n=4, X82107 n=3, X821-8 n=3, X82108 n=3, X831-4 n=4, X83104 n=4.

### Boosting PD and therapeutic potential of antimiRs via oleic acid conjugation

Previous results support that Ol conjugation improves the PD and therapeutic impact of the antimiR candidates X82107 and X82108 *in vivo*. We hypothesized that these benefits could be sequence independent and would also apply to other antimiRs. To test this hypothesis we designed miR-218 antimiRs, as this miRNA had previously been shown to upregulate MBNL1 in human cells and mouse models and could also have therapeutic potential in DM.^24,26,27^ This assessment involved a 5-day treatment of *HSA^LR^* mice with 3 mg/kg of an Ol-conjugated variant (X83104) compared to the unconjugated antimiR-218 control (X831-4) with an identical chemical structure (Figure 4A,B). The spider graphs in Figure 4C illustrate that Ol-conjugated antimiRs against miR-23b or miR-218 consistently outperformed their unconjugated counterparts across all evaluated parameters (data from FVB control mice are included in dataset S1). These conjugates showed greater efficiency in reducing target miRNA levels, indicating enhanced molecular targeting. Additionally, treatment with the conjugated antimiRs resulted in a significant increase in MBNL1 protein levels and *Mbnl1* and *Mbnl2* transcript expression, underscoring their ability to restore critical molecular markers associated with DM1. Splicing recovery, including improved *Mbnl1* exon 5 inclusion, was notably higher in mice treated with the conjugates, and conjugated variants also provided the most robust recovery of functional parameters, suggesting that the Ol modification enhances therapeutic potential. The overall performance of conjugated antimiRs highlights the advantages of Ol conjugation for improving molecular and functional outcomes in this disease model.

### Sustained therapeutic effects of conjugated oligos

To build on the results obtained for prospective lead candidates X82107 and X82108 from the 5-day single-dose experiments, we tested the activity duration of these 3’Ol-conjugated molecules after the same 3 mg/kg IV dose by evaluating tissue response at 5, 7, and 15 days post-injection (Figure 5A, dataset S1). Throughout the study period, both antimiRs exhibited a sustained increase in Mbnl1 protein levels compared to phosphate-buffered saline (PBS)-injected controls, with the peak effect observed at 5 days post-treatment (Figure 5B). Notably, X82108 consistently outperformed X82107 in this parameter at each time point. These findings are consistent with a significant overall PSR on days 5 and 7, during which protein levels were still substantially higher than in disease controls, and X82108 provided notably better splicing rescue than X82107 (Figure 5C, dataset S1). Both antimiRs demonstrated a comparable peak in normalized grip strength at 5 and 7 days post-injection, nearing normal values before declining to PBS-injected levels by day 15. In the context of myotonia, X82108 achieved a greater reduction than X82107 at day 5, with both converging at later time points (Figure 5D,E, dataset S1). Taken together, these results suggest that the activity duration for both lead candidates ranged between 7 and 15 days, with very similar outcomes observed for the two compounds.

**Figure 5.**
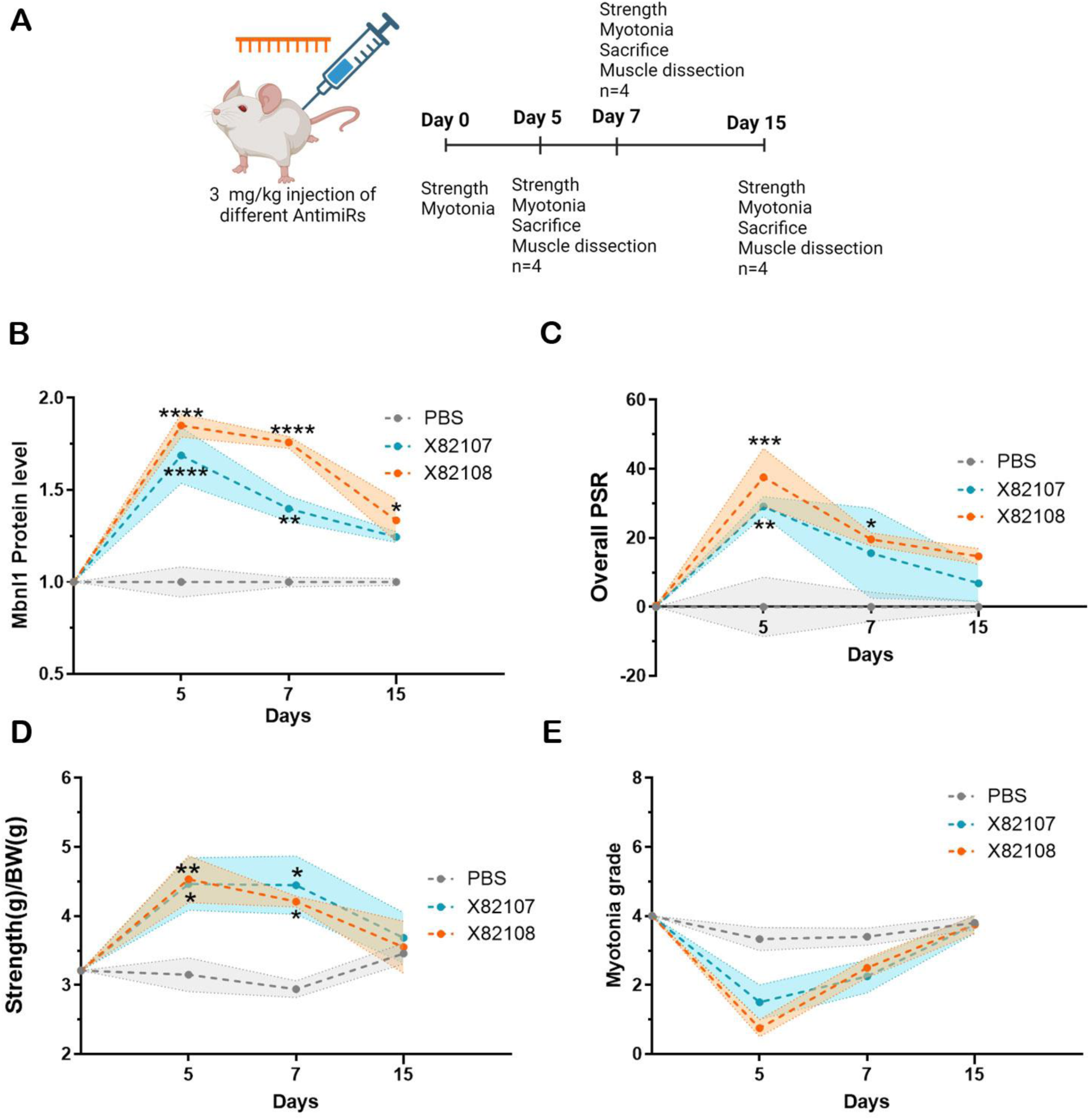
Selected antimiRs effects until day 15. **(A)** Schematic representation of the experimental design using a single 3 mg/kg IV injection in *HSA^LR^* mice. During 15 days after injection, several parameters were measured, and skeletal muscle tissues were collected for analysis: **(B)** MBNL1 levels of treated mice relative to saline controls and **(C)** overall PSR score in qd and gt muscles, **(D)** strength/body weight (g) and **(E)** myotonia grade at different time points after injection (AI). *: p<0.05; **: p<0.01; ***: p<0.001; ****: p<0.0001. The data were analyzed by one-way ANOVA or Kruskal-Wallis test if required, compared with untreated *HSA^LR^*mice (PBS). The shaded area indicates the error in each case.

### Dose-response efficacy of X82107 and X82108

Upon demonstrating the promising effects of Ol conjugation on X82107 and X82108 variants, we further investigated the efficacy of these molecules at a higher dose (6 mg/Kg), 15 days after injection, when the effects of the 3 mg/Kg injection were already substantially reduced. A single intravenous injection of 6 mg/kg of each variant was administered to *HSA^LR^* mice (Figure 6A,B, dataset S1), and several parameters were evaluated. Specifically, both antimiRs led to significantly higher levels of MBNL protein in the skeletal muscles qd and gt compared to PBS-treated animals. However, the increase observed with X82108 was notably higher than with X82107 (Figure 6C, dataset S1). This trend was also reflected in the reduction of miR-23b levels, which decreased by up to 80% in X82108-treated animals (dataset S1), and in a higher PSR achieved by this antimiR (Figure 6D, dataset S1). These molecular changes translated into functional improvements, with treated mice displaying similarly increased front leg grip strength, normalized to body weight (Figure 6E, dataset S1). Both candidates demonstrated a significant increase in PSR and normalized strength 15 days after injection at 6 mg/kg (Figure 6F), confirming prolonged activity compared to results at the lower dose of 3 mg/kg conjugated with oleic acid or cholesterol.^52^

**Figure 6.**
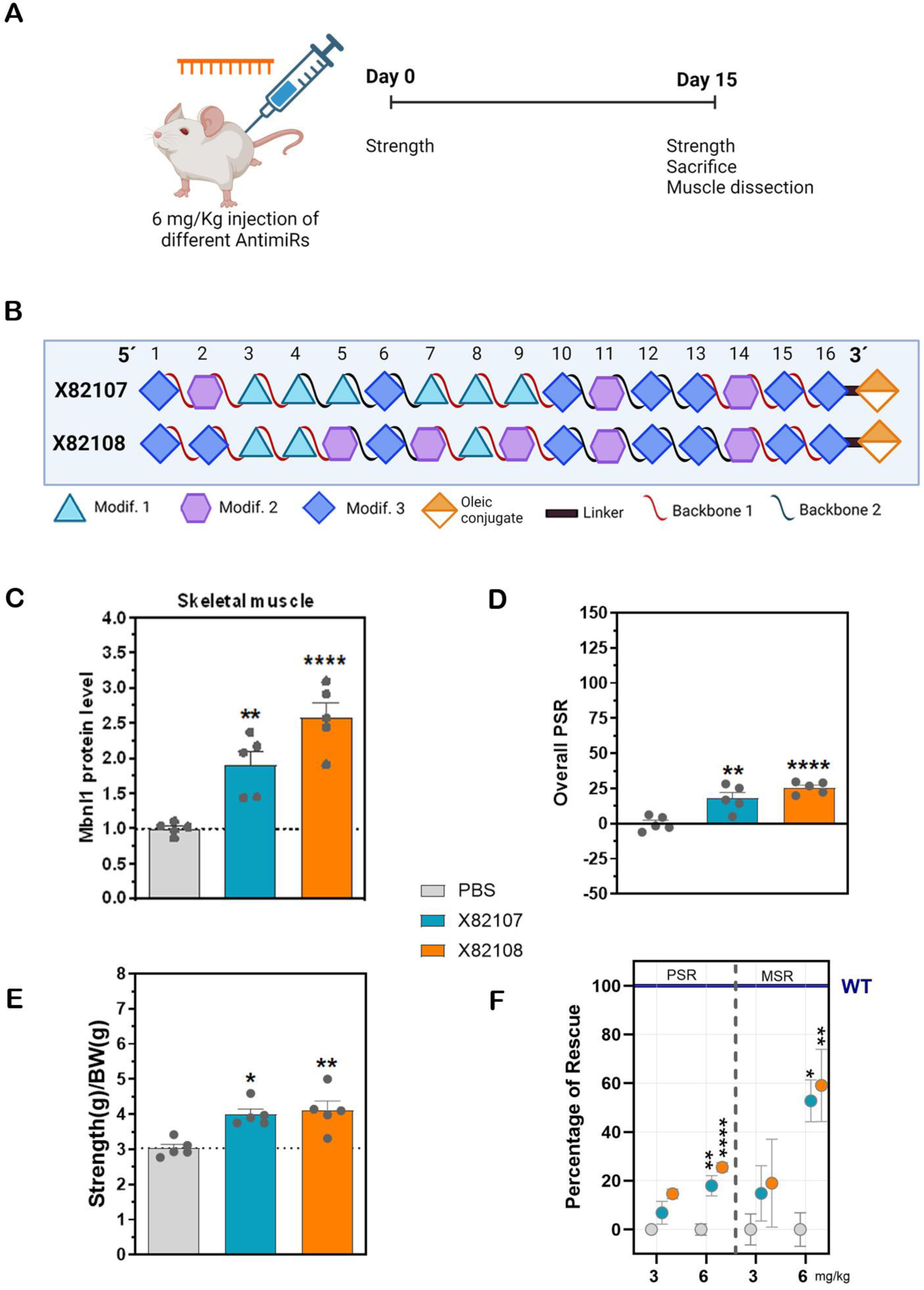
*In vivo* preclinical analysis with leading antimiR candidates. **(A)** Schematic representation of the experimental design using a single 6 mg/kg IV injection in *HSA^LR^* mice. **(B)** Schematic representation of the antimiRs tested. Modifications 1, 2, and 3 refer to 2’-O-Methyl RNA nucleotides, 2’-O-MOE RNA nucleotides, and LNA nucleotides. Backbones 1 and 2 represent a mix of PS (phosphorothioate) and PO (phosphodiester) linkages. **(C)** MBNL1 levels of treated mice relative to saline controls in both muscles (mean value of qd and gt). **(D)** Overall PSR scores. **(E)** Front leg grip strength normalized to body weight. **(F)** Percentage of PSR and MSR (muscle strength rescue) rescues. Blue line indicates wild-type levels. Error bars indicate mean ± SEM. *: *p*<0.05; **: *p*<0.01; ***: *p*<0.001; ****: *p*<0.0001. The data were analyzed by ANOVA one-way test or Kruskal-Wallis test, when necessary, compared with untreated *HSA^LR^* mice (PBS). Color legend: PBS: gray; X82107: blue; X82108: orange.

Overall, both molecules exhibited a dose-dependent duration of effect, further supporting their potential as promising therapeutic candidates.

### *In vitro* assessment of renal and immune activation profiles of X82107 and X82108

Potential renal toxicity was assessed using an *in vitro* model with RPTEC/TERT1 cells, human renal proximal tubular epithelial cells modified with telomerase reverse transcriptase for long-term studies. RPTEC/TERT1 cells are ideal for nephrotoxicity, renal metabolism, and drug absorption studies, as they retain primary renal cell functions, including electrolyte transport, toxin response, and epithelial barrier integrity;^70^ furthermore, EGFR activation is a pivotal mediator for renal fibrosis and has a major role in activating pathways that mediate podocyte injury and loss in diabetic nephropathy.^71^ After treatment with antimiRs, extracellular epidermal growth factor (EGF) levels rose significantly at the highest dose of 100 µM for antimiR X82108 compared to PBS-treated controls (Figure 7A), while a moderate but non-significant increase was observed for X82107 at the same dose (Figure 7B). No substantial changes in EGF levels were detected at 1, 10, or 30 µM for either compound. Parallel MTS assays revealed significant cytotoxicity at 100 µM for both compounds, with cell viability inhibition exceeding 80% (Figure 7C,D). This aligns with the accumulation of extracellular EGF, as non-viable cells have reduced EGF uptake capacity.

**Figure 7.**
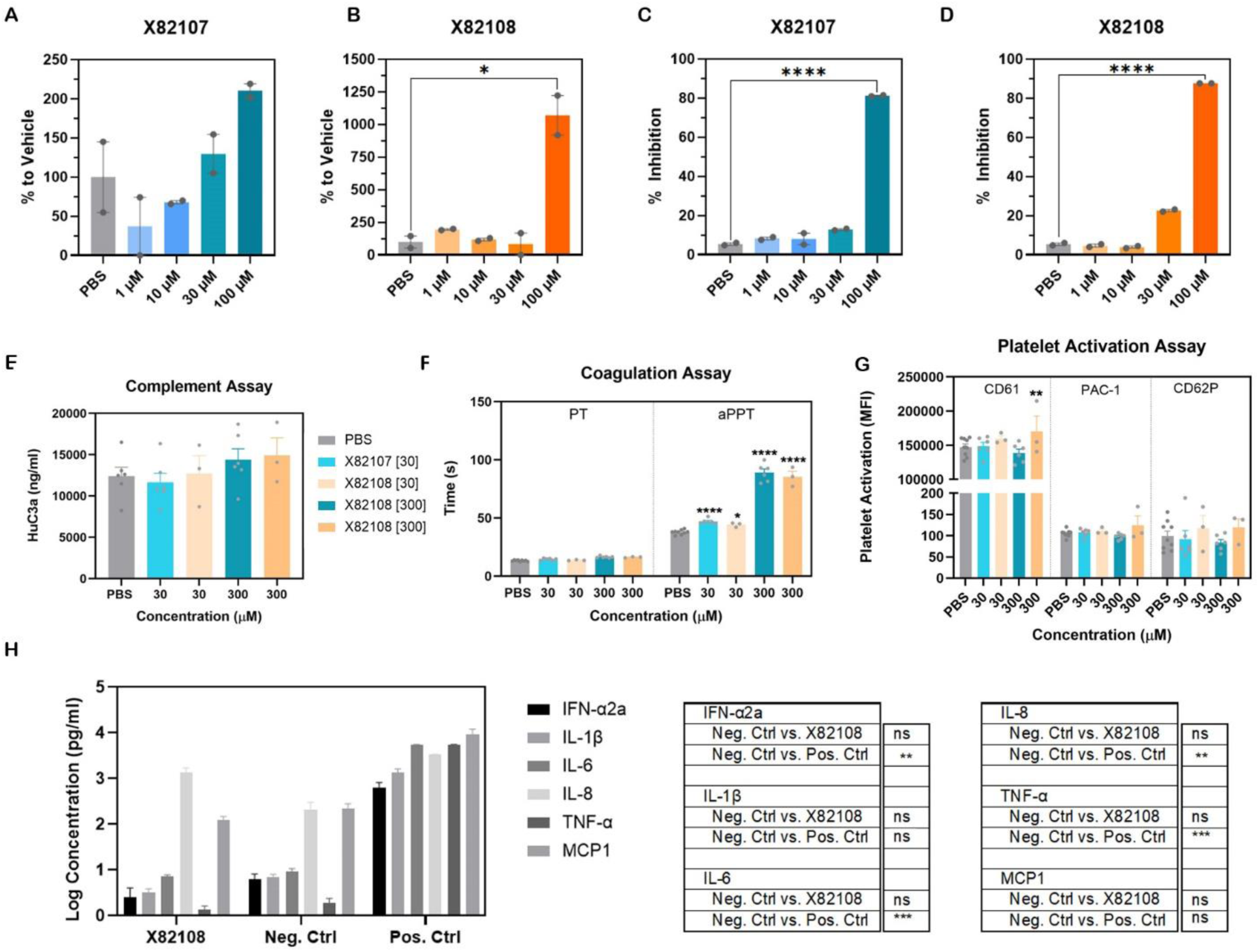
Toxicity and immune activation experiments with ascending doses of X82107 and X82108. EGF measurement **(A, B)** and cell proliferation inhibition assay **(C, D)** in RPTEC/TERT1 cells upon antimiR treatment. **(E-H)** Immune activation analysis in blood samples from three different donors: **(E)** activated complement (C3a) measurement, **(F)** coagulation assay (prothrombin and activated partial thromboplastin times), and **(G)** total and activated platelet count as measured by several cell surface markers. **(H)** Cytokine release in human peripheral blood mononuclear cells (PBMCs) upon X82108 treatment. *: *p*<0.05; **: *p*<0.01; ***: *p*<0.001; ****: *p*<0.0001 according to one-way ANOVA or Kruskal-Wallis test if required, compared with negative control. aPPT: activated partial thromboplastin time; CD61: integrin beta-3; CD62P: platelet surface P-selectin; IFN: interferon; IL: interleukin; MCP1: monocyte chemoattractant protein-1; PAC1: activated GP IIb/IIIa; PBS: phosphate buffered saline; PT: prothrombin time; TNF: tumor necrosis factor.

In summary, significant effects on extracellular EGF levels and cell viability were observed only at 100 µM, a concentration typically associated with toxicity in these assays. At lower doses, both antimiRs displayed a favorable safety profile.

We also conducted *in vitro* tests in human blood to assess the potential immunogenic modulation and safety of lead candidates X82107 and X82108. Neither compound activated the complement system, as evidenced by the unchanged C3a levels observed in serum and plasma samples, even at a high dose of 300 µM (Figure 7E). Given that different chemical modifications in oligonucleotide-based drugs can significantly alter the pharmacokinetic and toxicological profiles of antisense ON, including potential effects on coagulation times,^72^ we measured these parameters in normal human plasma. In coagulation assays, both X82107 and X82108 produced a significantly prolonged activated partial thromboplastin time (aPTT, which measures the intrinsic coagulation pathway) at 30 µM and higher doses in normal human plasma, while the prothrombin time (PT, extrinsic pathway) remained unchanged for both molecules compared to the PBS control (Figure 7F). Platelet count and activation levels remained similar to control across all doses for both compounds (Figure 7G). An additional assessment of cytokine release in human peripheral blood mononuclear cells (PBMCs) showed no significant cytokine increase following X82108 treatment (Figure 7H). Overall, our preclinical data support a favorable toxicological profile for lead compounds X82107 and X82108, with no significant safety-related red flags.

### AntimiR-23b-mediated MBNL upregulation is conserved in different mammal species

Human, rat (*Rattus norvegicus*) and pig (*Sus scrofa*) mature miR-23b sequences are identical. Given this conservation, we tested the hypothesis that our lead antimiRs could similarly antagonize miR-23b function upon transfection and upregulate MBNL1 protein expression in cell lines from these species. Transfection of X82108 in fibroblasts of these two animals led to a significant increase in *Mbnl1* levels and a dramatic reduction in miR-23b detection (Figure 8A,B). Importantly, these results confirm that miR-23b inhibition produces *Mbnl1* upregulation in different relevant preclinical mammalian species.

**Figure 8.**
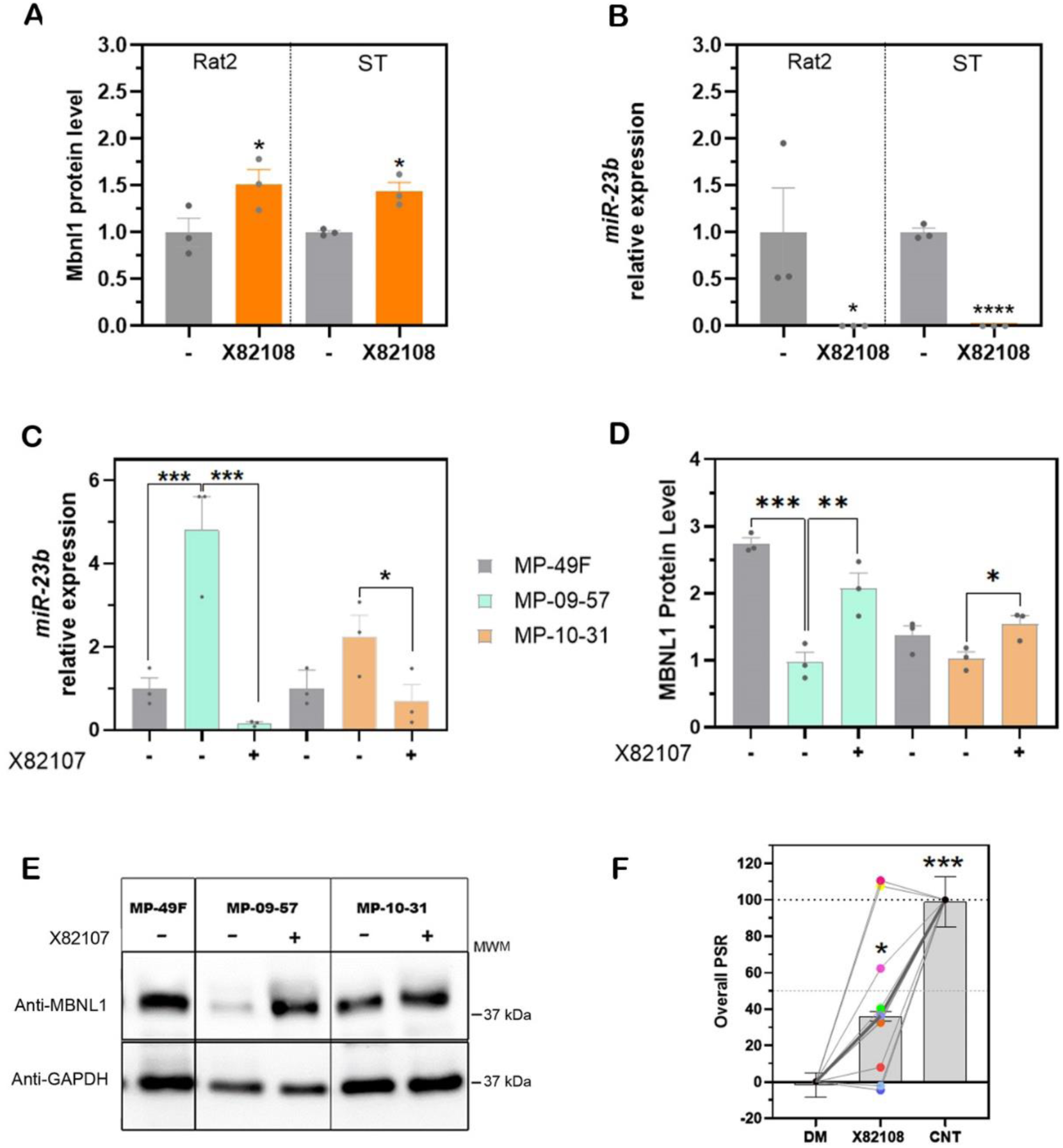
Confirmation of conserved antimiR-23b regulation activity on MBNL levels in various cell types. **(A)** MBNL1 levels and **(B)** miR-23b relative levels in Rat2 and ST cells upon treatment with X82108. *: *p*<0.05, ****: *p*<0.0001 according to Student’s t-test compared with non-treated cells; ST: *Sus scrofa* (testis). **(C)** miR-23b relative levels and **(D,E)** MBNL1 levels in primary myoblasts derived from DM1 patients upon treatment. MP-49F: control cell line; MP-09-57: DM1 cell line carrying 702 CTG repeats in *DMPK*; MP-10-31: DM1 cell line carrying 117 repeats. **(F)** Overall Percent Splicing Rescue (PSR) values for the adapted biomarker panel based on the Composite Alternative Splicing Index (CASI-22) in immortalized DM1 myotubes treated with 200 nM antimiR X82108; each colored dot represents an individual splicing event. *ANK2*-green; *ATP2A1*-fuchsia; *OPA1*-bright green; *BIN1*-light blue, *DMD*-yellow; *GFPT1*-red; *GOLGA4*- Blue; *MBNL1*-Orange, *INSR*-light violet; *NFIX*- mustard yellow and *SOS1*- magenta. -: untreated; +: treated with X82107; *: p<0.05; **: p<0.01; ***: p<0.001. The data were analyzed by one-way ANOVA or Kruskal-Wallis test if required, compared with untreated control cell line (MP-49F). ST: *Sus scrofa* (testis).

To rule out cell line bias, X82107 and X82108 were tested on patient-derived primary myoblasts after 10 days of differentiation. This has already been reported for X82108 ^27^ and is shown here for X82107 in two cell lines from that same study (Figure 8C–E). Briefly, treatment of primary human myoblast cell lines MP09-57 (carrying 702 CTG repeats in *DMPK*) and MP10-31 (carrying 117 repeats) with X82107 resulted in a marked reduction of miR-23b detection (Figure 8C). Overexpression of miR-23b and reduction in the presence of lead antimiRs correlated well with a significant increase of MBNL1 protein levels that approached or surpassed those observed in healthy control cells (MP-49F; Figure 8D,E). Results from Figure 8 and ^27^ demonstrate that lead antimiRs can significantly upregulate MBNL1 levels in diverse genetic backgrounds and CTG repeat expansion sizes. Additionally, treatment with antimiR-23b significantly reduced the number of foci in these cell lines at different time points, as well as increased MBNL1 protein in a diffused, free-form state in the nucleus. ^27^

### Transcriptomic analysis of miR-23b inhibition: miR-23b targets analysis and splicing correction

To further characterize the molecular consequences of miR-23b inhibition, we evaluated transcriptomic alterations in immortalized DM1 myotubes, where miR-23b is markedly upregulated (Figure S4), consistent with observations in *HSA^LR^* mice ^25^ and primary patient-derived myotubes (Figure 8C and ^27^). Myotubes were treated with 200 nM antimiR X82108 for 3 days and subjected to total RNA-seq. Sylamer analysis ^73^ was performed to determine whether miR-23b seed–complementary sequences were enriched or depleted among differentially expressed genes (DEGs), highlighting the miRNA-driven regulatory effects. Moreover, we additionally evaluated the impact of miR-23b inhibition on the transcriptomic biomarker Composite Alternative Splicing Index (CASI-22), a robust clinical biomarker consisting of 22 DM1-specific splicing events strongly correlated with muscle function and widely used to evaluate disease-modifying treatments ^62^.

Genome-wide Sylamer analysis revealed a significant overrepresentation of the miR-23b-3p seed motif (AATGTGA) among the downregulated mRNAs in DM1 compared with controls (Figure S5A and Table S5), consistent with the repressive activity of elevated miR-23b levels. Following antimiR treatment, this enrichment was found only among upregulated genes (Figure S5B), indicating selective derepression of endogenous miR-23b targets. These results confirm that miR-23b targets are repressed in the disease condition and antimiR X82108 specifically blocks miR-23b activity, leading to increased expression of its repressed targets in the DM1 cellular context.

Given that RNA mis-splicing is the core molecular hallmark of DM1, we assessed whether miR-23b inhibition could ameliorate splicing defects using the CASI-22 biomarker panel. Of the 17 CASI events detectable in the cells, 11 were significantly altered, 3 exhibited borderline significant changes, and 3 remained unchanged relative to controls. Notably, antimiR X82108 rescued the overall PSR up to 35% and significantly improved 80% of the splicing events altered, with 20% of them fully rescued (Figure S5C, Figure 8F). Many of the corrected events correspond to splicing alterations closely associated with muscle-strength decline in DM1—most notably *ATP2A1*, *DMD*, *NFIX*, *OPA1*, and the autoregulatory exon of *MBNL1*—underscoring their functional and clinical relevance. ^62^

Collectively, these transcriptomic and splicing analyses demonstrate that antimiR X82108 exerts a precise and therapeutically meaningful molecular effect, derepressing miR-23b targets, and improving key DM1 biomarkers implicated in muscle dysfunction.

## DISCUSSION

We aimed to enhance the therapeutic potential of miR-23b inhibition in DM1 by developing optimized antimiRs. Inhibiting miR-23b has been shown to boost MBNL1 levels and counteract critical DM1 molecular defects. Given its elevated expression in DM1,^27^ we hypothesized that reducing miR-23b activity could mitigate disease effects. Starting with the non-optimized antimiR-23b molecule X820 (antagomiR-23b),^52^ we employed a two-step *in vitro* and *in vivo* screening strategy to refine miR-23b-targeting sequences, chemical modifications, and lipid conjugations, optimizing activity and toxicity in DM1 muscle cells. This approach combined the best features from each screening, yielding advanced candidates. Animal experiments confirmed the target engagement of these optimized molecules, validating the use of cell-based assays to streamline *in vivo* testing.^74^

The PK/PD characteristics of ON drugs are strongly influenced by their chemical modifications and specific properties of any associated conjugates. ^67,75^ To be effective in DM1, therapy must reach skeletal muscles, which can constitute up to approximately 40% of the total body mass in average healthy adults. However, the first clinical trial evaluating a therapeutic ON targeting *DMPK* transcripts in DM1 was discontinued due to insufficient drug levels in muscle tissue, even at the highest doses tested.^44^ This underscores the inherent delivery challenges faced in treating genetic myopathies. Most ONs approved for human use are administered systemically by intravenous or subcutaneous injections to achieve broad distribution throughout the body. ON plasma concentration peaks immediately after administration, followed by rapid distribution to major body tissues within hours, except for the central nervous system.^47^

Receptor-specific delivery has been addressed by conjugating *DMPK*-degrading ONs to monoclonal antibodies or Fab fragments recognizing the transferrin receptor, aimed at enhancing muscle tissue uptake.^35^ The mechanism by which Ol conjugation may confer desirable properties to an ON has not been formally elucidated, but an intriguing hypothesis is that lipids can facilitate either fusion with or disruption of the endosomal membrane, allowing the antisense ON to escape into the cytoplasm.^76,77^ ONs conjugated with fatty acids longer than C16 are known to form self-assembled vesicular structures, which may further enhance their intracellular delivery. The interaction of these fatty acid chains with mouse and human serum albumin (MSA/HSA) results in the formation of stable adducts, with a near-linear correlation between fatty acid-ON hydrophobicity and binding strength to albumin.^67,68^ This enhanced albumin binding likely prolongs circulation time, improves systemic stability, and facilitates efficient transport across endothelial barriers to reach target tissues, including skeletal muscle. These properties may collectively contribute to the superior therapeutic potential observed with Ol-conjugated antimiRs. Importantly, while we compared unconjugated and conjugated ONs at the same dose in mg/kg, the higher molecular weight of the conjugated ONs means that fewer molecules were administered relative to their unconjugated counterparts. As a result, the observed increase in potency may be slightly underestimated, further underscoring the enhanced efficacy of the Ol-conjugated ONs.

Safety is the primary concern in early drug development for human use, so we prioritized ON candidates that demonstrated the greatest therapeutic window. This prioritization strategy enabled efficient progression from initial hits to lead candidates. Interestingly, significant toxicity was observed only at very high doses of antimiRs *in vitro* (e.g., 30–100 µM) for most of the compounds tested. This yielded remarkable (over 100-fold) differences between the TC_50_ and the EC_50_ concentrations.

Our results indicate that lead compounds X82107 and X82108 do not activate human platelets or increase C3a in serum, with X82108 showing a slight, non-clinically meaningful prolongation in coagulation time at high doses. Overall, the compounds exhibit a safe toxicological profile, without adverse effects on coagulation or platelet activation. Although microRNAs regulate numerous biological pathways, their inhibition does not necessarily lead to widespread systemic effects. Functional knockout and CRISPR/Cas9 studies have shown that deletion of miR-23b or the miR-23b/27b/24-1 cluster in mice produces only mild phenotypes, such as a minor increase in pain sensitivity in neuropathic models or altered glucose homeostasis under a high-fat diet, without major developmental or systemic defects. ^78,79^ These findings, together with reports identifying miR-23b inhibition as therapeutically beneficial in cancer, support a favorable safety profile for miR-23b inhibition. ^80^

Since miRNAs are short and highly conserved regulatory molecules, the same antimiR molecule can usually be tested in several mammalian animal models during regulatory preclinical steps.^81^ In this case, after obtaining solid results in mice, we moved on to test our leading candidates in normal rat and pig fibroblasts. This paves the way for future use of minipigs as a large animal model in preclinical toxicology testing of antimiRs, given the numerous physiological similarities with humans, especially in cardiac studies.^82^ Notably, these experiments confirmed that the antimiR mechanism of action was conserved across mammalian species, even without expressing toxic CTG repeat expansions. Finally, the robust increase in MBNL1 levels in primary myoblast derived from two DM1 patients upon X82107 or X82108^27^ treatment demonstrated the effectiveness of these antimiRs across different human genetic backgrounds affected by DM1. Moreover, the new transcriptomic analyses further support the specificity and therapeutic relevance of miR-23b inhibition, showing selective derepression of endogenous targets and partial correction of key DM1-associated splicing defects, thereby complementing our previously published transcriptomic study in eight primary DM1 myotube lines treated with antimiR-23b ^27^.

Studies with the original antagomiR-23b molecule determined an optimal dose of 12.5 mg/kg to significantly increase MBNL1 levels *in vivo.*^24^ In comparison, the new lead candidates exhibited lower toxicity and higher potency, achieving the same effect on MBNL1 levels 5 days after administration with an approximately fourfold lower dose (3 mg/kg) and sustaining this level of MBNL1 upregulation for at least 15 days with approximately half the dose (6 mg/kg). Notably, this enhanced potency also translated into improved alternative splicing correction, as reflected by a significantly higher overall PSR at both 3 mg/kg and 6 mg/kg doses, with the strongest splicing rescue observed at the peak of MBNL1 upregulation (Figure 5B,C).

While our study demonstrates the efficacy and safety of X82107 and X82108 in preclinical models, several limitations must be considered. First, although we observed robust MBNL1 upregulation and splicing correction, further studies are needed to evaluate whether these molecular improvements translate into sustained functional benefits beyond muscle strength and myotonia reduction. Additionally, while *in vivo* biodistribution analyses confirmed efficient skeletal muscle uptake, the extent of off-target effects in other tissues was not fully characterized by these experiments. Given the high conservation of miR-23b across species, the potential for unintended modulation of physiological pathways outside DM1-affected tissues was carefully examined in additional studies. Another key limitation is the absence of long-term toxicity data from these experiments, particularly regarding immune activation and hepatic or renal clearance after repeated dosing, which require further investigation to dose humans on a chronic basis.

To bridge the gap toward clinical application, the next steps should be to include large-animal studies to confirm the efficacy and pharmacokinetics of these lead compounds in models with greater physiological similarity to humans, such as minipigs and non-human primates. Additionally, regulatory preclinical studies require detailed dose-escalation experiments to establish the maximum tolerated dose and long-term safety, particularly in the context of systemic administration. As ON-based therapies for neuromuscular diseases continue to evolve, complementary formulation strategies may further enhance the bioavailability and T_index_ of these compounds. Ultimately, clinical trial design will incorporate biomarker-driven endpoints, including MBNL1 target engagement and splicing correction, to facilitate a rigorous evaluation of therapeutic efficacy in DM1 patients.

In conclusion, our approach combining *in vitro* screening, rational design, and *in vivo* validation efficiently identifies potent and safe candidates. This strategy enhances therapeutic development in the antimiR field and advances DM1 treatments by optimizing delivery and reducing toxicity.

## Acknowledgments

We thank Inmaculada Noguera for veterinary assistance at the University of Valencia SCSIE animal facility. Part of the equipment employed in this work has been funded by Generalitat Valenciana and co-financed with ERDF funds (OP ERDF of Comunitat Valenciana 2014-2020). Antibody MB1a (4A8) was provided by the MDA Monoclonal Antibody Resources. This work was supported by “la Caixa” Banking Foundation grant HR17-00268 (RA, AL-M, GG), Generalitat Valenciana grants PROMETEO/2020/081 and CIPROM/2023/22 (RA), Instituto de Salud Carlos III grant DTS19/0128 (RA), Generalitat Valenciana predoctoral grant FDEGENT/2020/011 (IG-M), Torres Quevedo post-doctoral fellowship PTQ2020-011110 (EC-H), CDTI NEOTEC grant SNEO-20201136 (BL), GVA-IVACE grant IMIDTA/2021/65 (BL), Talent Promotion Program-Line 3 of GVA-AVI grant INNTA3/2023/16 (DP-L), and Instituto de Salud Carlos III grant PI21/00557 (NN-G and AL-M). Illustrations were created with BioRender.

## Author contributions

Conceptualization: RA, EC-H, BL.

Methodology: IG-M, MAV, NM, MC-S, NB, AD-M.

Validation: EC-H, AG-R, DP-L, NB.

Formal analysis: IG-M, EC-H, MC-S, AG-R, DP-L, AC-R, MCH, AD-M.

Investigation: IG-M, EC-H, MC-S, AG-R, DP-L, AC-R, MCH, AH-L, AG-B, NM, MD.

Resources: RA, NN-G, AL-M, GG.

Visualization: IG-M, EC-H, AD-M.

Funding acquisition: RA, BL, AL-M, GG.

Project administration: RA, BL, AL-C.

Supervision: RA, EC-H, GG, AL-M.

Writing – original draft: RA, AL-C.

Writing – review & editing: RA, IG-M, EC-H, AL-C.

## Data and code availability

All materials used in this study are available upon reasonable request. AntimiRs will require a material transfer agreement. All data are available in the main text or the supplementary materials. RNA-seq data are publicly available with accession number S-BSST2306 in the BioStudies database.

