## Supplemental information for "Enhanced muscle uptake of chemically optimized miR-23b antisense oligonucleotides as lead compounds for Myotonic Dystrophy type 1"

### Supplemental figures and tables

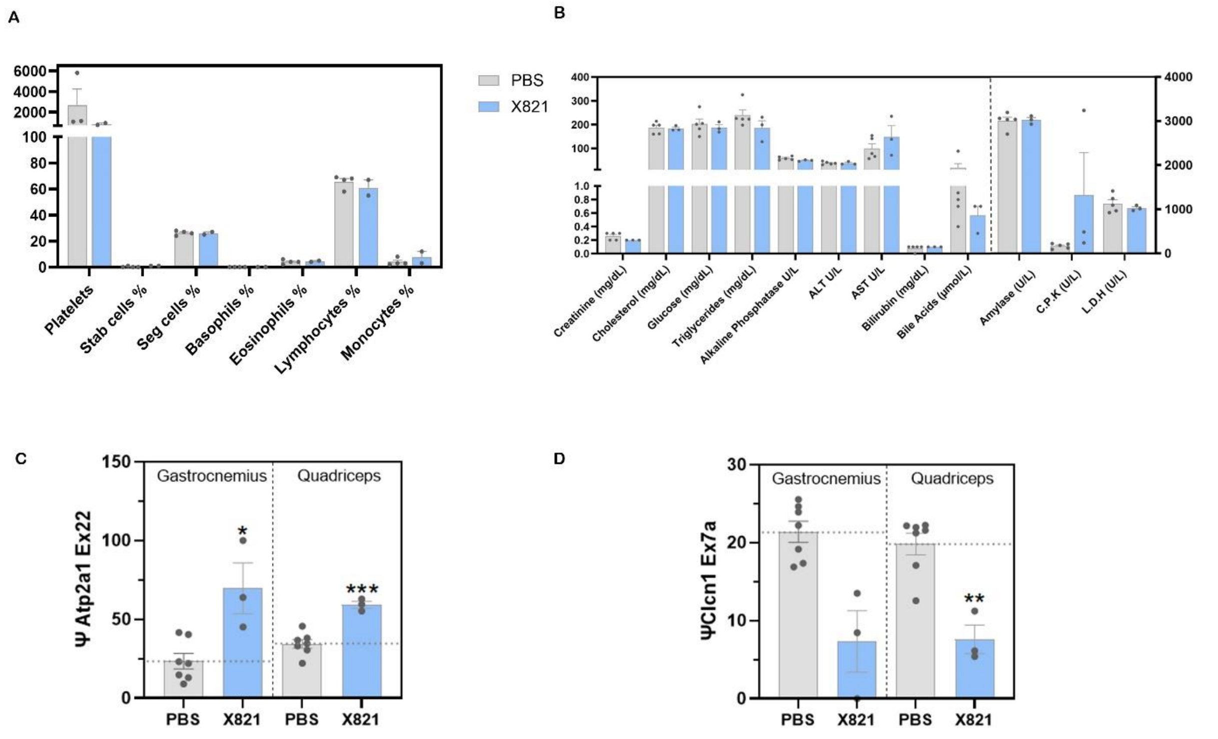


**Figure S1. Further safety and efficacy measurements of X821 treatment at 100 mg/kg in *HSA$^{LR}$* mice. (A)** White blood cell differential count and **(B)** blood serum biochemistry were measured in total blood extracted before sacrifice. **(C,D)** Quantification of the percentage of exon inclusion (Ψ) of the indicated exons and muscles. Error bars indicate mean ± SEM. *: p<0.05; **: p<0.01; ***: p<0.001. The data were analyzed by one-way ANOVA or Kruskal-Wallis test if required, compared with untreated DM1 mice (PBS). PBS n=7, X821 n=3.

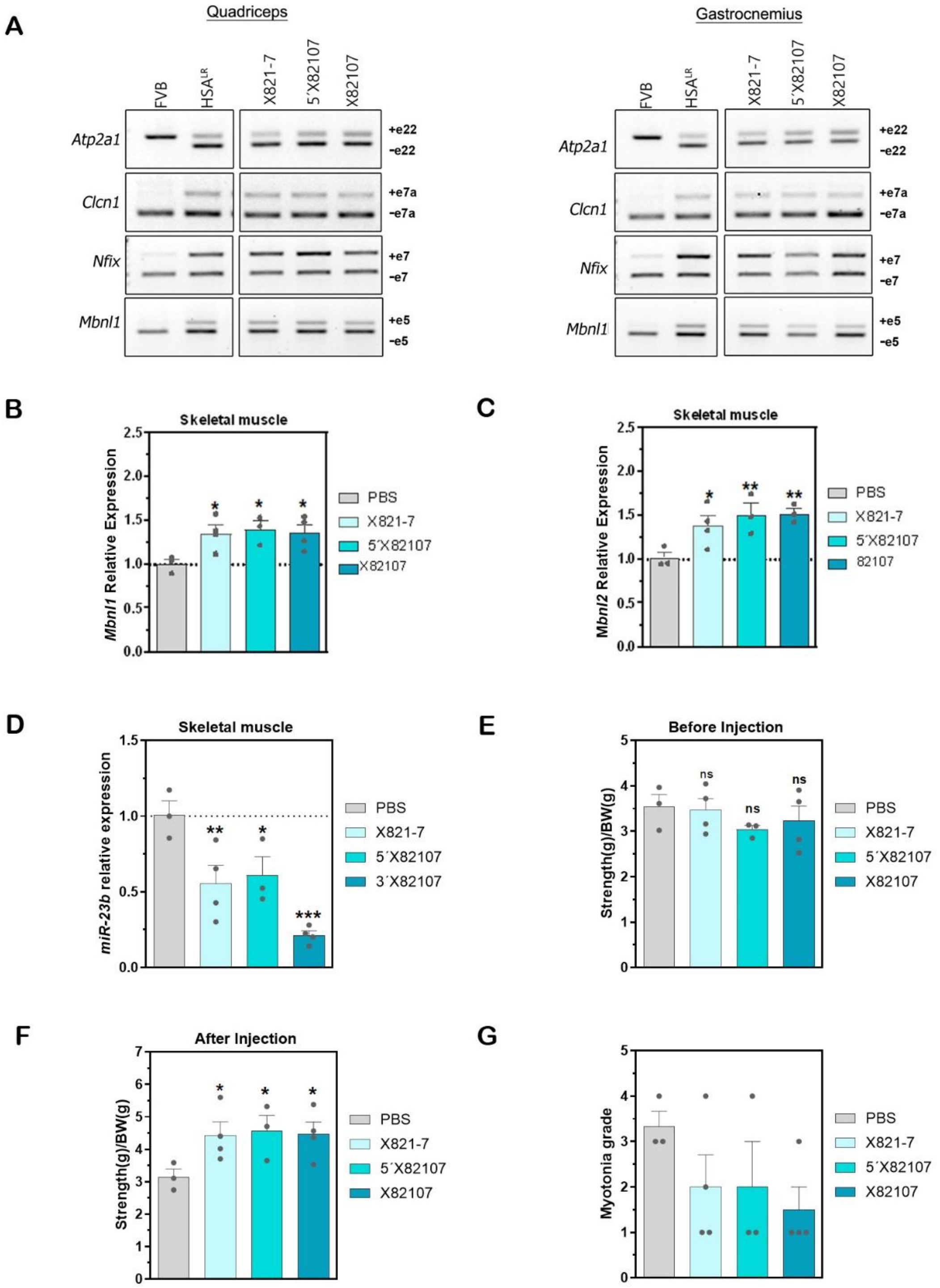


**Figure S2. Single-dose experiment with leading antimiR derivatives from the *in vitro* screening.** Selected compounds were tested *in vivo* using a single 3 mg/kg IV injection in *HSA*$^{LR}$ mice. After five days, several parameters were measured, and skeletal muscle tissues were dissected for analysis. Splice recovery of *Atp2a1* ex22, *Clcn1* ex7a, *Nfix* ex7, and *Mbnl1* ex5 in **(A)** quadriceps and gastrocnemius. **(B)** MBNL1, **(C)** MBNL2 and **(D)** miR-23b relative expression were also quantified after treatment. Functional recovery

[**(E–F)** strength and **(G)** myotonia] was measured **(E)** before and **(F,G)** after injection with the different tested compounds. Error bars indicate mean ± SEM. *: $p<0.05$; **: $p<0.01$; ***: $p<0.001$. The data were analyzed by one-way ANOVA or Kruskal-Wallis test if required, compared with untreated $HSA^{LR}$ mice (PBS). Individual values are indicated as data points. PBS n=3, X821-7 n=4, 5'X82107 n=3, X82107 n=4.

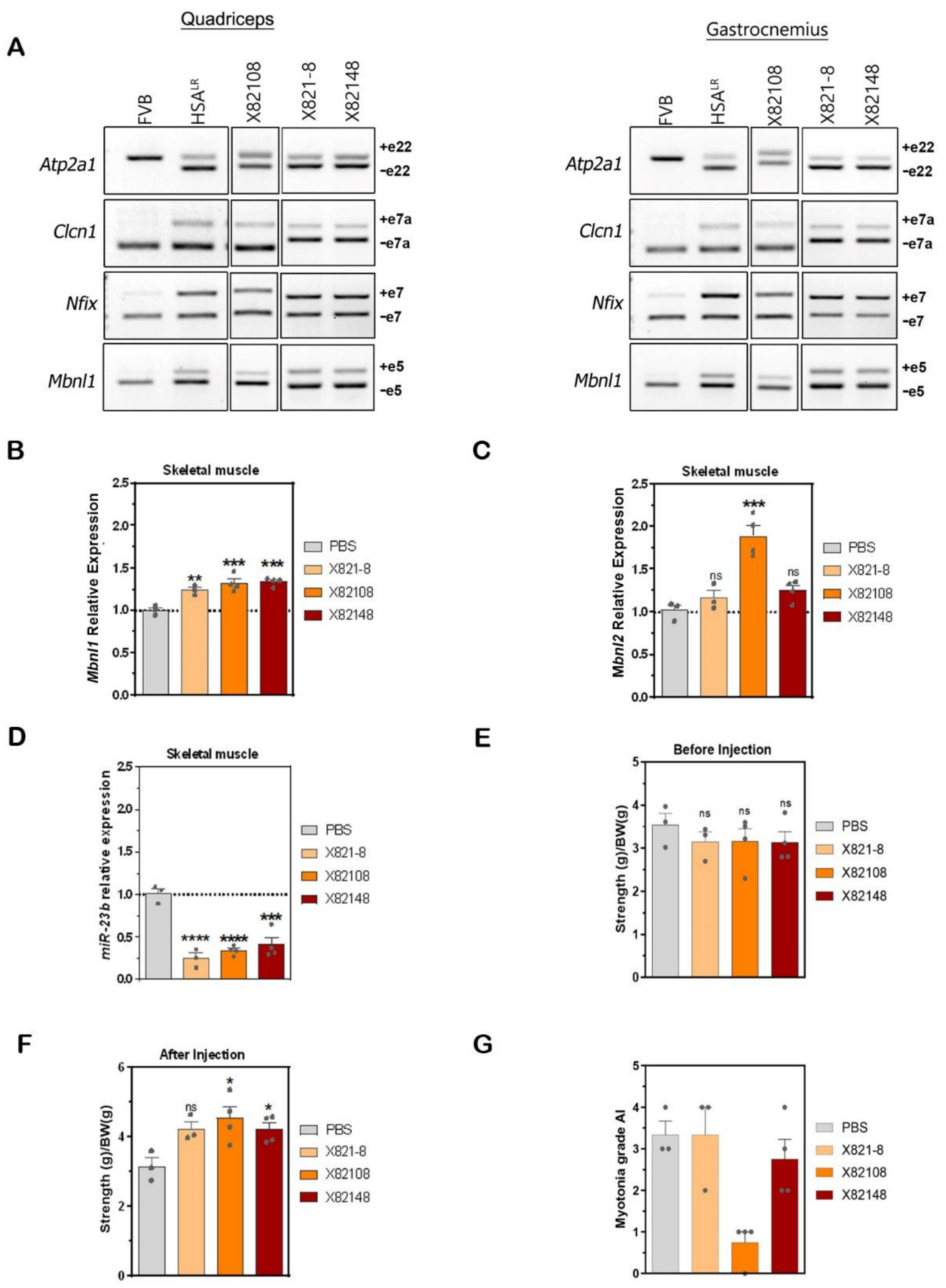


**Figure S3. Single-dose experiment with leading antimiR derivatives from the previous experiment.** Selected compounds were tested *in vivo* using a single 3 mg/kg IV injection in HSA$^{LR}$ mice. After 5 days, several parameters were measured, and skeletal muscle tissues were dissected for analysis. Splice recovery of *Atp2a1* ex22, *Clcn1* ex7a, *Nfix* ex7, and *Mbnl1* ex5 in **(A)** quadriceps and gastrocnemius. **(B)** MBNL1, **(C)** MBNL2 and **(D)** miR-23b relative expression were also quantified after treatment. Functional recovery [**(E–F)** strength and **(G)** myotonia] was measured **(E)** before and **(F,G)** after injection with the different tested compounds. Error bars indicate mean ± SEM. *: p<0.05; **: p<0.01; ***: p<0.001; ****: p<0.0001. The data were analyzed by one-way ANOVA or Kruskal-Wallis test if required, compared with untreated *HSA*$^{LR}$ mice (PBS). Individual values are indicated as data points. PBS n=3, X821-8 n=3, X82108 n=4, X82148 n=4.

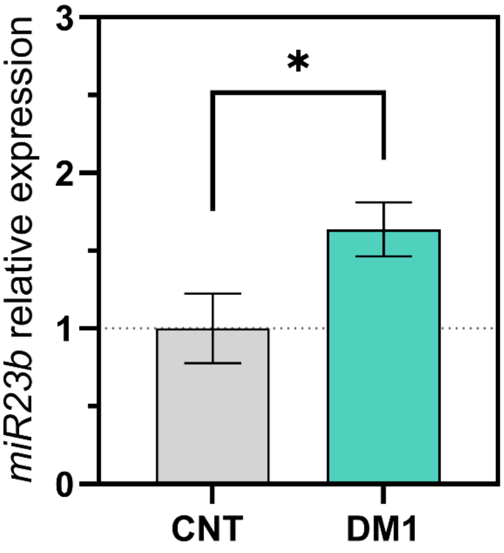


**Figure S4. miR-23b is upregulated in DM1 myotubes.** miR-23b expression was quantified after 3 days of differentiation by RT-qPCR. Error bars indicate mean ± SEM. *: p<0.05; **: p<0.01; ***: p<0.001; ****: p<0.0001. The data were analyzed by unpaired t test, compared with control (CNT). CNT n=5, DM1 n=7.

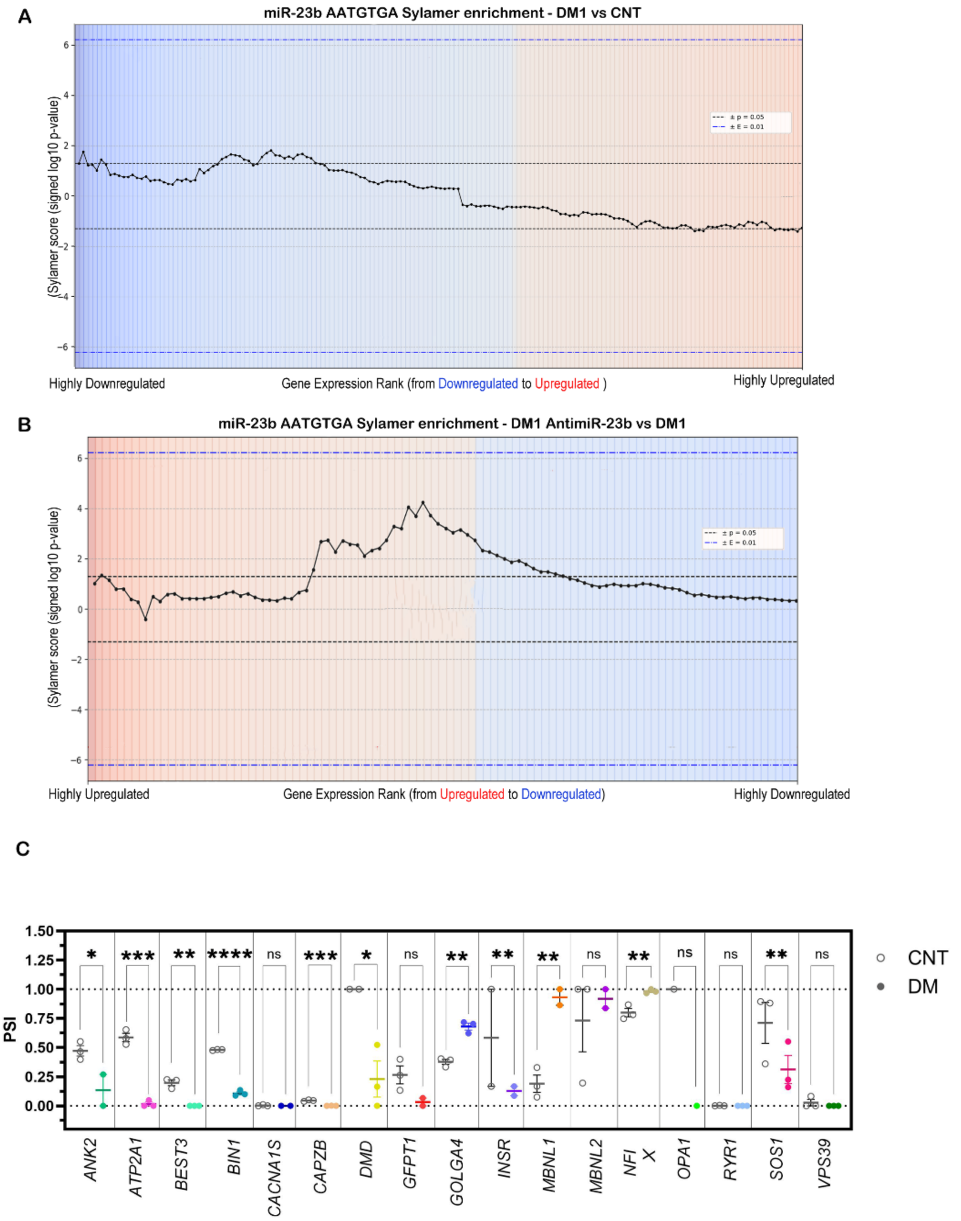


**Figure S5. Sylamer enrichment landscape plots for miR-23b seed sequences in DM1 myotubes.** Sylamer enrichment plots showing the distribution of the miR-23b-3p seed word (AATGTGA) along ranked gene lists for **(A)** DM1 versus control and **(B)** DM1 treated with 200 nM antimiR-23b versus DM1. The x-axis represents genes ordered from most downregulated to most upregulated in (**A**) or from most upregulated to most downregulated in (**B**). The y-axis shows the signed −log10(P-value) of enrichment or depletion for the seed motif across cumulative bins. Positive values indicate enrichment, and negative values indicate depletion. Dashed horizontal lines correspond to significance thresholds: black ($p < 0.05$) and blue ($p < 0.01$). **(C)** Percent spliced-in (PSI) values for individual DM1-relevant alternative splicing events included in the adapted CASI-22 biomarker panel, measured in immortalized control (CNT) and DM1 myotubes. For the DM1 condition, each splicing event is represented with a different color. Each dot corresponds to an individual biological replicate. Statistical significance was assessed by one-way ANOVA or Kruskal–Wallis test as appropriate.

**Table S1. Screening of antimiR candidates: chemistry and sequences.** This table indicates the sequence for each oligonucleotide tested in DM1 TDMs. The dose range tested always included five levels, but lower concentrations were required for TC and EC determinations of certain compounds. The following parameters were calculated: $EC_{50}$, $TC_{50}$, $E_{max}$ (maximum fold change), and $T_{index}$ (therapeutic index). The gray rows show the results of a compound identical to the original antimiR-23b (20, 21) but from a provider that used a different linker between the oligonucleotide and the cholesterol moiety (X820), and an identical antimiR lacking cholesterol, which were used as our benchmarks. Sequence key: N (A, G, T, C): DNA and RNA nucleotides and (C): 5-methyl-2'-O-methyl cytidine.

| Name | Sequence (5' > 3') | $TC_{50}$ | $EC_{50}$ | $E_{max}$ | $T_{index}$ |
|---|---|---|---|---|---|
| X820 | GGUAAUCCCUGGCAAUGUGAU-CHOLESTEROL | 0.857 | 0.204 | 7.548 | **31.7** |
| X82-- | GGUAAUCCCUGGCAAUGUGAU | 1.704 | 0.264 | 1.478 | **9.5** |
| X821 | AT(C)(C)(C)TGG(C)AATGTGA | 14.000 | 0.010 | 2.870 | **4018.3** |
| X822 | C(C)(C)TGGCAATGUGAT | 0.471 | 0.002 | 2.916 | **686.7** |
| X823 | C(C)(C)TGG(C)AATGTGAT | 0.025 | 0.001 | 1.655 | **51.7** |
| X824 | AT(C)(C)(C)TGG(C)AATGTGAT | 1.510 | 0.030 | 1.763 | **88.7** |
| X825 | TAATCCCTGGCAATGTGAT | 2.271 | 0.068 | 2.247 | **75.0** |
| X826 | CCCTGGCAATGTGAT | 0.598 | 0.019 | 1.758 | **55.3** |
| X827 | ATCCCTGGCAATGTGAT | 0.263 | 0.019 | 2.343 | **32.4** |
| X828 | TAAT(C)(C)(C)TGGCAATGTGAT | 1.604 | 0.235 | 3.564 | **24.3** |
| X829 | ATCCCTGGCAATGTGA | 0.234 | 0.058 | 4.817 | **19.4** |
| X8210 | CCCUGGCAAUGUGAT | 2.850 | 0.010 | 1.570 | **447.4** |
| X8211 | TAAT(C)C(C)TGG(C)AATGTGA | 0.128 | 0.021 | 2.497 | **15.2** |
| X8212 | ATCCCTGGCAATGTGA | 0.275 | 0.040 | 1.918 | **13.2** |
| X8213 | C(C)(C)UGGCAAUGUGAT | 1.640 | 0.043 | 4.194 | **159.9** |
| X8214 | ATCCCTGGCAATGTGAT | 0.117 | 0.026 | 2.191 | **9.9** |
| X8215 | GGUAAUC(C)(C)UGGCAAUGUGAT | 0.400 | 0.051 | 4.924 | **38.6** |
| X8216 | AATCCCTGGCAATGTGAT | 0.481 | 0.113 | 2.701 | **11.5** |
| X8217 | AATCCCTGGCAATGTGAT | 0.370 | 0.357 | 4.078 | **4.2** |
| X8218 | ATCCCTGGCAATGTGAT | 0.175 | 0.110 | 1.299 | **2.1** |
| X8219 | TAAT(C)C(C)TGG(C)AATGTGA | 0.110 | 0.102 | 1.473 | **1.6** |
| X8220 | GGUAAUCCCUGGCAAUGUGAT | 5.562 | 0.255 | 2.153 | **47.0** |

**Table S2. Screening of antimiR conjugate candidates.** The sequence for each oligonucleotide used in DM1 TDMs is indicated. The following parameters were calculated: $EC_{50}$, $TC_{50}$, $E_{max}$ (maximum fold change), and $T_{index}$ (therapeutic index). The gray row shows the results for our benchmark antimiR-23b (35, 36). Sequence key: N (A, G, T, C): DNA and RNA nucleotides and (C): 5-methyl-2'-O-methyl cytidine.

| **Name** | **Sequence (5' > 3')** | **$TC_{50}$** | **$EC_{50}$** | **$E_{max}$** | **$T_{index}$** |
|---|---|---|---|---|---|
| X820 | GGUAAUCCCUGGCAAUGUGAU-CHOL | 0.857 | 0.204 | 7.548 | 31.70 |
| X82-- | GGUAAUCCCUGGCAAUGUGAU | 1.704 | 0.264 | 1.478 | **9.53** |
| X8200 | OLEYL-GGUAAUCCCUGGCAAUGUGAU | 4.040 | 0.010 | 2.726 | **1101.40** |
| X8201 | LINOLEYL-GGUAAUCCCUGGCAAUGUGAU | 40.630 | 0.186 | 1.835 | **400.94** |
| X8202 | TOCOPHEROL-GGUAAUCCCUGGCAAUGUGAU | 3.460 | 0.100 | 2.211 | **76.59** |
| X8203 | CHOLESTEROL-GGUAAUCCCUGGCAAUGUGAU | 1.027 | 1.279 | 13.654 | **10.96** |
| X8204 | PALMITOYL-GGUAAUCCCUGGCAAUGUGAU | 2.156 | 0.460 | 1.775 | **8.32** |
| X8205 | ELAIDOYL-GGUAAUCCCUGGCAAUGUGAU | 0.700 | 1.341 | 1.628 | **0.85** |
| X8206 | STEAROYL -GGUAAUCCCUGGCAAUGUGAU | 0.432 | 0.730 | 1.848 | **1.09** |

**Table S3. Leading antimiR candidates selected for further preclinical testing**. Sequence key: N (A, G, T, C): DNA and RNA nucleotides and (C): 5-methyl-2'-O-methyl cytidine. The conjugate used in each case is indicated in the table.

| **Name** | **Sequence (5' > 3')** |
|---|---|
| X821-7 | AT(C)(C)(C)TGG(C)AATGTGA |
| 5'X82107 | OLEYL-AT(C)(C)(C)TGG(C)AATGTGA |
| X82107 | AT(C)(C)(C)TGG(C)AATGTGA-OLEYL |
| X821-8 | AT(C)(C)CTGGCAATGTGA |
| X82108 | AT(C)(C)CTGGCAATGTGA-OLEYL |
| X82148 | AT(C)(C)CTGGCAATGTGA-PALITOYL |
| X831-4 | TTAGATCAAGCACAA |
| X83104 | TTAGATCAAGCACAA-OLEYL |

**Table S4. Probes and primers used for RT-qPCR.** Sequence key: 6FAM: Fluorescein modification; IAbRQSp: Iowa Black RQ-Sp; MAX: NHS ester modification; BHQ_1: Black Hole Quencher-1.

| Gene | | Sequence (5'→3') |
|---|---|---|
| *Mbnl1* (mouse) | **Probe** | /6FAM/TCGCAAATCAGCTGTGAGGAGATTCCCT/IAbRQSp/ |
| | **Primer Forward** | TACCGATTGCACCACCAAAC |
| | **Primer Reverse** | GCTGCTTTCAGCAAAGTTGTC |
| *Mbnl2* (mouse) | **Probe** | /6FAM/CCCGGCAGACAGCACCATGATCGA/IAbRQSp/ |
| | **Primer Forward** | GAGACAGACTGCCGCTTTG |
| | **Primer Reverse** | GGTTACGGTGTTGTCGTTTGT |
| *Gapdh* | **Probe** | /MAX/-CGCCTGGTCACCAGGGCTGCT-/BHQ_1/ |
| | **Primer Forward** | CAACGGATTTGGTCGTATTGG |
| | **Primer Reverse** | TGATGGCAACAATATCCACTTTACC |

**Table S5. Identifiers of the splicing events analyzed in adapted CASI-22 biomarker panel.** The table provides, for each analyzed splicing event, the gene symbol, gene name, chromosomal location, strand orientation, and genomic start and end coordinates.

| Gene Symbol | Gene Name | Chrom | Strand | Start | End |
|---|---|---|---|---|---|
| *ANK2* | ankyrin 2 | NC_000004.12 | + | 113372533 | 113372625 |
| *ATP2A1* | ATPase sarcoplasmic/endoplasmic reticulum Ca2+ transporting 1 | NC_000016.10 | + | 28903700 | 28903741 |
| *BEST3* | bestrophin 3 | NC_000012.12 | - | 69694370 | 69694464 |
| *BIN1* | bridging integrator 1 | NC_000002.12 | - | 127060596 | 127060640 |
| *CACNA1S* | calcium voltage-gated channel subunit alpha1 S | NC_000001.11 | - | 201054505 | 201054561 |
| *CAPZB* | capping actin protein of muscle Z-line subunit beta | NC_000001.11 | - | 19342752 | 19342864 |
| *DMD* | dystrophin | NC_000023.11 | - | 31126642 | 31126673 |
| *GFPT1* | glutamine--fructose-6-phosphate transaminase 1 | NC_000002.12 | - | 69354259 | 69354312 |
| *GOLGA4* | golgin A4 | NC_000003.12 | + | 37361243 | 37361305 |
| *INSR* | insulin receptor | NC_000019.10 | - | 7150497 | 7150532 |
| *MBNL1* | muscleblind like splicing regulator 1 | NC_000003.12 | + | 152446704 | 152446757 |
| *MBNL2* | muscleblind like splicing regulator 2 | NC_000013.11 | + | 97366459 | 97366553 |
| *NFIX* | nuclear factor I X | NC_000019.10 | + | 13078613 | 13078735 |
| *OPA1* | OPA1 mitochondrial dynamin like GTPase | NC_000003.12 | + | 193617784 | 193617837 |
| *RYR1* | ryanodine receptor 1 | NC_000019.10 | + | 38523915 | 38523929 |
| *SOS1* | SOS Ras/Rac guanine nucleotide exchange factor 1 | NC_000002.12 | - | 38989270 | 38989314 |
| *VPS39* | VPS39 subunit of HOPS complex | NC_000015.10 | - | 42192066 | 42192098 |

### Supplemental material and methods

*Cell proliferation assay*

DM1 and control fibroblasts seeded at $10^5$ cells/ml in 96-well plates were transfected 24 h later with antimiRs, as previously described; after 96 h, cell proliferation was measured using the CellTiter 96® Aqueous Non-Radioactive Cell Proliferation Assay (Promega, Madison, Wisconsin). The absorbance levels were determined using an Infinite M200 PRO plate reader (Tecan, Männedorf, Switzerland). RPTEC/TERT1 cells were seeded into 96-well plates (Greiner Bio-One™, 655180), at 20,000 cell/well density, and treated as previously explained. Cell viability was determined using the same procedure as fibroblasts upon medium harvesting on day 6.

*Protein extraction, quantitative dot blot (QDB) and western blot (WB) assays*

For the activity assay, cells were seeded in 6-well plates at a density of $8x10^4$ cells per well and transfected 24 h later with antimiRs, as described in the previous sections. For total protein extraction, human muscle cells were sonicated while mouse muscles (gastrocnemius, gt and quadriceps, qd) were homogenized in Pierce® RIPA buffer (Thermo Scientific, Waltham, Massachusetts) supplemented with protease and phosphatase inhibitor cocktails (Roche Applied Science, Penzberg, Germany). Total protein was quantified with a Pierce® BCA protein assay kit (Thermo Scientific, Waltham, Massachusetts) using bovine serum albumin as standard. For the immunodetection assay, 1 μg/well of cell and 2 μg/well of mice samples were denatured (100°C for 5 min) and loaded in QDB plates (Quanticision Diagnostics Inc, Research Triangle Park, North Carolina). Quantifications by QDB were performed as described in [1,2]. Quantifications of Mbnl1 protein levels by western blot were as described in [3].
